# Building alternative consensus trees and supertrees using *k*-means and Robinson and Foulds distance

**DOI:** 10.1101/2021.03.24.436812

**Authors:** Nadia Tahiri, Bernard Fichet, Vladimir Makarenkov

## Abstract

Each gene has its own evolutionary history which can substantially differ from the evolutionary histories of other genes. For example, some individual genes or operons can be affected by specific horizontal gene transfer and recombination events. Thus, the evolutionary history of each gene should be represented by its own phylogenetic tree which may display different evolutionary patterns from the species tree that accounts for the main patterns of vertical descent. The output of traditional consensus tree or supertree inference methods is a unique consensus tree or supertree. Here, we describe a new efficient method for inferring multiple alternative consensus trees and supertrees to best represent the most important evolutionary patterns of a given set of phylogenetic trees (i.e. additive trees or *X*-trees). We show how a specific version of the popular *k*-means clustering algorithm, based on some interesting properties of the Robinson and Foulds topological distance, can be used to partition a given set of trees into one (when the data are homogeneous) or multiple (when the data are heterogeneous) cluster(s) of trees. We adapt the popular Caliński-Harabasz, Silhouette, Ball and Hall, and Gap cluster validity indices to tree clustering with *k*-means. A special attention is paid to the relevant but very challenging problem of inferring alternative supertrees, built from phylogenies constructed for different, but mutually overlapping, sets of taxa. The use of the Euclidean approximation in the objective function of the method makes it faster than the existing tree clustering techniques, and thus perfectly suitable for the analysis of large genomic datasets. In this study, we apply it to discover alternative supertrees characterizing the main patterns of evolution of SARS-CoV-2 and the related betacoronaviruses.

## INTRODUCTION

Most of the conventional consensus and supertree inference methods generate one candidate tree for a given set of input gene phylogenies. However, the topologies of gene phylogenies can be substantially different from each other due to possible horizontal gene transfer, hybridization or intragenic/intergenic recombination events by which the evolution of the related genes could be affected (Bapteste et al. 2004). Each gene phylogeny depicts a unique evolutionary history which does not always coincides with the main patterns of vertical descent depicted by the species tree (Szöllősi et al. 2014). In order to infer a reliable species phylogeny, the related gene trees should be merged, while minimizing topological conflicts presented in them (Maddison et al. 2007). In this context, two scenarios can be envisaged: First, trees to be merged are constructed for the same set of taxa, which are usually associated with tree leaves (i.e. the case of *consensus trees*), and second, trees to be merged are constructed for different, but mutually overlapping, sets of taxa (the case of *supertrees*).

A large variety of methods have been proposed to address the problem of reconciliation of multiple trees defined on the same set of leaves in order to infer a consensus tree. The most known types of consensus trees are the strict consensus tree, the majority-rule consensus tree, and the extended majority-rule consensus tree (Day and McMorris 2003; Bryant 2003; Felsenstein 2004). However, in practical evolutionary studies, we rarely deal with phylogenies defined on the same set of taxa, and thus the consensus tree inference problem is transformed into the supertree inference problem (Bininda-Emonds 2004). Several approaches have been proposed to synthesize collections of small phylogenetic trees with partial taxon overlap into comprehensive supertrees including all taxa found in the input trees (Wilkinson et al. 2007). The most known of them are Strict Supertree (Sanderson et al. 1998), Matrix Representation with Parsimony (MRP) (Baum 1992; Ragan 1992; Bininda-Emonds 2003), Parsimony Supermatrix (Driskell et al. 2004; Ciccarelli et al. 2006), Majority-Rule supertrees (Cotton and Wilkinson 2007), Maximum Likelihood (ML) supertrees (Steel and Rodrigo 2008), SuperFine (Swenson et al. 2011), Multi Level Supertrees (MLS) (Berry et al. 2012), and Subtree Prune-and-Regraft (SPR) distance-based supertrees (Whidden et al. 2014). The MRP methods proceed by a matrix-like aggregation of separately inferred partial trees. The trees derived from these independent analyses are then combined to produce a single MRP matrix used to reconstruct the supertree of all sources of taxa (de Queiroz and Gatesy 2007). In the parsimony supermatrix methods all systematic characters are integrated into a single phylogenetic matrix which is used to analyze all characters simultaneously in order to build a supermatrix tree. The strict supertree includes the bipartitions (or splits) that agree with all bipartitions present in the input phylogenies, while the rest of the tree consists of unresolved sub-trees. The concept of strict and loose supertrees has been well described by McMorris and Wilkinson (2011), who showed that these types of supertrees are natural generalizations of the corresponding consensus trees.

The implementation of the famous “Tree of Life” (ToL) project intended to infer the largest possible species phylogeny became feasible due to collaborative efforts of biologists and nature enthusiasts from around the world (Maddison et al. 2007). The approach adopted by the project organizers consists of gradual partitioning of the complex tree reconstruction problem into several sub-problems, followed by merging the obtained sub-trees. Indeed, such an approach produces thousands of small trees which should be assembled to create ToL. Precisely, the problem can be viewed as twofold: First, we have to infer small sub-trees of ToL (i.e. often gene trees representing different evolutionary histories), the most commonly defined on different, by mutually overlapping, sets of taxa, and second, we have to merge these small sub-trees into *one* or *several supertrees* using a supertree reconstruction method that allows combining trees inferred for different sets of taxa (Szöllősi and Daubin 2012). In this context, the application of a method that infers *multiple supertrees* (i.e. a supertree clustering method) would help discover and regroup plausible alternative evolutionary scenarios for several sub-trees of ToL.

Unfortunately, most of the traditional consensus tree and supertree inference methods return as output a single consensus tree or supertree. Thus, in many instances, these methods are not informative enough, as they do not preserve alternative evolutionary scenarios that characterize sub-groups of genes that have undergone similar reticulate evolutionary events (e.g. genes whose evolution has been affected by similar horizontal gene transfers). Maddison (1991) was the first to formulate the idea of multiple consensus trees describing the Phylogenetic Islands method. He observed that the consensus trees of the islands can differ substantially from each other, and that they are usually much better resolved than the consensus tree of the whole set of taxa. The most intuitive approach for discovering and regrouping genes sharing similar evolutionary histories is clustering their gene phylogenies. In this context, Stockham et al. (2002) proposed a tree clustering algorithm based on *k*-means (Lloyd 1957; MacQueen 1967; Bock 2007) and the quadratic version of the Robinson and Foulds topological distance (Robinson and Foulds 1981). The clustering algorithm proposed by Stockham et al. aims at inferring a set of strict consensus trees that minimizes the information loss. It proceeds by determining the consensus trees for each set of clusters in all intermediate partitioning solutions tested by *k*-means. This makes the algorithm of Stockham et al. very expensive in terms of the running time. Bonnard et al. (2006) introduced a method, called MultiPolar Consensus (MPC), which produces a minimum number of consensus trees displaying all splits of a given set of phylogenies whose support is above a predefined threshold. Guénoche (2013) proposed the Multiple Consensus Trees (MCT) method for partitioning a group of phylogenetic trees into one or several clusters. The method of Guénoche computes a generalized partition score to determine the most appropriate number of clusters for a given set of gene trees. Finally, Tahiri et al. (2018) have recently proposed a fast tree clustering method based on *k*-medoids. To the best of our knowledge, all the methods proposed for building multiple alternative phylogenies assume that the input trees have identical sets of taxa, thus working in the consensus tree context. Therefore, the relevant and challenging problem of inferring multiple supertrees still needs to be addressed appropriately. In this paper, we describe a new method that can be used to infer both alternative consensus trees and supertrees. We present some interesting properties of the Robinson and Foulds topological distance and show that it should not be used in its quadratic form in tree clustering. We adapt the popular Ball and Hall (Ball-Hall 1967), Caliński-Harabasz (Caliński and Harabasz 1974), Gap (Tibshirani et al. 2001), and Silhouette (Rousseeuw 1987) cluster validity indices to *k-*means tree clustering. Our method is validated through a comprehensive simulation study. It is then applied to discover alternative supertrees characterizing the main patterns of evolution of SARS-CoV-2 and the related betacoronaviruses (Lam et al. 2020).

## MATERIALS AND METHODS

A phylogenetic tree is an unrooted leaf-labeled tree in which each internal node, representing an ancestor of some contemporary species (i.e. taxa), has at least two children and all leaves, representing contemporary species, have different labels. Our method takes as input a set Π of *N* phylogenetic trees defined on the same (or different, but mutually overlapping) set(s) of leaves and returns as output one or several disjoint clusters of trees from which the corresponding consensus trees (or supertrees) can then be inferred. Our approach relies of the use of a fast version of the popular *k*-means algorithm adapted for tree clustering. We define and compare different variants of the *k*-means objective function suitable for clustering trees.

*K*-means (Lloyd 1957; MacQueen 1967) is an unsupervised data partitioning algorithm which iteratively regroups *N* given objects (i.e. phylogenetic trees in our case) into *K* disjoint clusters. The content of each cluster is chosen to minimize the sum of intracluster distances. The most commonly used distances in the framework of *k*-means are the Euclidean distance, the Manhattan distance, and the Minkowski distance (Bock 2007). The problem of finding an optimal partitioning according to the *k*-means least-squares criterion is known to be NP-hard (Mahajan et al. 2009). This fact has motivated the development of a number of polynomial - time heuristics, most of them having the time complexity of *O*(*KNIM*), for finding an approximate clustering solution, where *I* is the number of iterations in the *k*-means algorithm and *M* is the number of variables characterizing each of the *N* objects. In this work, we used the version of *k*-means implemented by Makarenkov and Legendre in the program OVW (Makarenkov and Legendre 2001).

The Robinson and Foulds distance (*RF*), also known as the symmetric-difference distance (Dong et al. 2010), between two trees is a well-known metric widely used in computational biology to compare phylogenetic trees defined on the same set of taxa (Robinson and Foulds 1981; Makarenkov and Leclerc 2000; Bordewich et al. 2009). The *RF* distance is a topological distance. It does not take into account the length of the tree edges. As shown by Barthélemy and McMorris (1986), the majority-rule consensus tree of a set of trees is a median tree of this set in the sense of the *RF* distance. This fact justifies the use of this distance in tree clustering.

One of the main advantages of our method is that the proposed version of *k*-means does not recompute the consensus trees or supertrees for all intermediate clusters of trees, but estimates the quality of each intermediate tree clustering using approximation formulas based on the properties of majority-rule consensus trees and the *RF* distance. This allows for a much faster clustering of a given set of input phylogenies without compromising on the quality of the resulting consensus trees or supertrees.

### Approximation by the Euclidean distance

The traditional *k*-means algorithm (MacQueen 1967) partitions a given dataset into *K* (*K* > 1) disjoint clusters according to its objective function based on a specific distance (e.g. Euclidean or Minkowski distance) and the selected cluster validity index (e.g. Caliński-Harabasz, Silhouette or Dann index). Most of the traditional cluster validity indices take into consideration both intragroup and intergroup cluster evaluations. However, we cannot use the standard objective functions or cluster validity indices when clustering trees. Here, we discuss the main modifications that should be introduced into the conventional *k*-means algorithm in order to adapt it to tree clustering.

In case of tree clustering, the objective function of the method can be defined as follows:

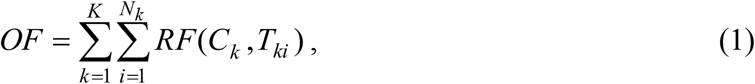

where *K* is the number of tree clusters, *N*_*k*_ is the number of trees in cluster *k, RF*(*C*_*k*_, *T*_*ki*_) is the *RF* distance between tree *i* of cluster *k*, denoted *T*_*ki*_, and the majority-rule consensus tree (any other type of consensus trees could be considered here) of cluster *k*, denoted *C_k_*.

Still, the computation of the majority-rule, or of the extended majority-rule, consensus tree is time-consuming. The running time of the method computing any of these consensus trees is *O*(*n*^2^ + *nN*^2^) (Wareham 1985), where *n* is the number of leaves in each tree and *N* is the number of trees. Even though an optimal algorithm for computing the majority-rule consensus tree, running in *O*(*nN*), has been proposed by Jansson et al. (2013), this time complexity cannot be achieved when the trees are defined by their Newick strings, as it is typically done in evolutionary biology (Felsenstein 2013). In the algorithm of Jansson et al., the trees are defined by their sets of clusters and its space consumption is *O*(*n*^2^*N*). Thus, the time complexity of a straightforward tree partitioning algorithm, such as that of Stockham et al. (2002), which recomputes the consensus trees after each *basic k-means operation* consisting in relocating an object (i.e. tree) from one cluster to another and then in reassessing the value of the objective function (Equation 1), is *O*((*n*^2^ + *nN* ^2^)*KI*), or *O*(*n* ^2^*N* + *nNKI*) when the trees are defined by their lists of clusters (here, *O*(*n*^2^*N*) operations are needed to transform *N* Newick strings into a list of *N* tree clusters), where *I* is the number of iterations in *k*-means. Tahiri et al. (2018) have recently proposed a *k*-medoids-based tree clustering method having the running time of *O*(*nN* ^2^ + *K* (*N* − *K*)^2^*I*), where *O*(*nN*^2^) is the time needed to precalculate the matrix of pair-wise *RF* distances of size (*N*×*N*) between all trees in Π.

In order to speed up the tree clustering process, we propose to use the following objective function *OF*_*EA*_ (Equation 2), which can be viewed as a Euclidean approximation of the objective function *OF* defined in Equation 1:

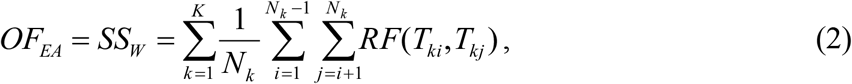

Where *SS*_*W*_ is the index of intragroup evaluation. Here, the *RF* distance, or more precisely its square root, replaces the traditional Euclidean distance used in *k*-means (see also Equation 17 in Appendix A.2 that provides two equivalent expressions for *SS*_*W*_ in case of the traditional Euclidean distance). The main advantage of using the objective function *OF*_*EA*_ is that we should not recompute the consensus tree for any intermediate cluster of trees considered by *k*-means, and that an object relocation operation consisting in finding the best cluster for a given tree *T* belonging to cluster *C* (i.e. when we try to relocate *T* into each of the *K*-1 clusters that are different from *C*) can be performed in *O*(*K*) time. Indeed, all pairwise *RF* distances between trees in a given set of input trees Π can be precalculated, and the sums of the *RF* distances between a given tree *T* and all elements of any tree cluster can first be precalculated and then updated at each step of *k*-means. The distance precalculation step can be completed in *O*(*nN*^2^) (Makarenkov and Leclerc 2000; Sul and Williams 2008). This leads to the total running time of *O*(*nN* ^2^ + *NKI*) for our extended majority-rule *k*-means based on Equation 2.

It is important to note that the Robinson and Foulds distance itself is not a Euclidean distance, but its square root is Euclidean. The proof of these important properties of the *RF* distance is presented in Appendix (A.1). Moreover, in Appendix (A.2) we explain how the popular Caliński-Harabasz (*CH*), Silhouette (*SH*), Ball and Hall (*BH*), and Gap cluster validity indices can be adapted to tree clustering with *k*-means.

### Approximation by the lower and the upper bounds, and by their mean

As discussed above, the Euclidean objective function *OF*_*EA*_ (Equation 2) is only an approximation of the objective functions *OF* defined in Equation 1. For instance, the consensus tree of the four trees presented in Figure A1 (see Appendix A.1) is a star tree, and the value of the objective function *OF* for them is (2 + 2 + 2 + 2) = 8, whereas the value of *OF*_*EA*_ is (2 + 2 + 4 + 4 + 2 + 2)/8 = 4.5. Thus, it would be interesting to find the lower and the upper bounds of the objective function *OF*, and use them in the clustering procedure. Theorem 1 below allows us to establish these bounds.

#### Theorem 1

*For a given cluster k containing N*_*k*_ *phylogenetic trees (i*.*e. additive trees or X-trees) the following inequalities hold*:

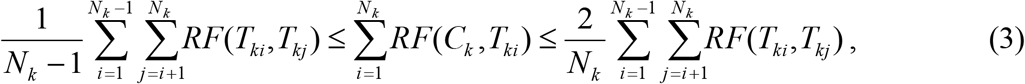

*where N*_*k*_ *is the number of trees in cluster k, T* _*ki*_*and T*_*kj*_ *are, respectively, trees i and j in cluster k, and C*_*k*_ *is the majority-rule consensus tree of cluster k*.

The proof of Theorem 1 is presented in Appendix A.2. It is worth noting that the lower and the upper bounds of 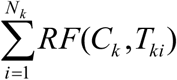 defined in Theorem 1 are identical when *N*_*k*_ = 2. Moreover, the value of the objective function defined by the upper bound of (3) divided by 2 (i.e. the function *OF*_*EA*_ defined in Equation 2) is smaller than the value of the objective function *OF* (Equation 1). Thus, according to the terminology of approximation theory, the criterion *OF*_*EA*_ is a factor-2 approximation of the criterion *OF*.

We can use these bounds as well as the middle of the interval defined by them, which is 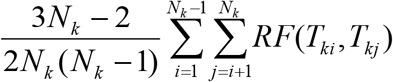, as an approximation of the contribution of cluster *k* to the *k*-means objective function *OF* defined by Equation 1.

For instance, the following objective function based on the middle of the interval established in Theorem 1 can be used as an approximation of the original objective function *OF*:

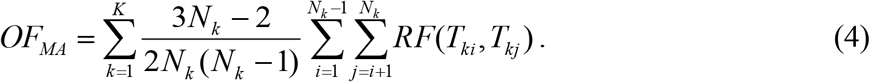

In a similar way, we can define the objective function based on the lower bound of *OF*:

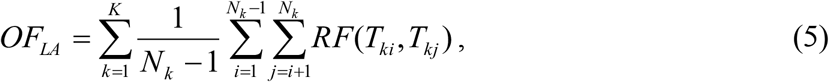

and the upper bounds of *OF* as well:

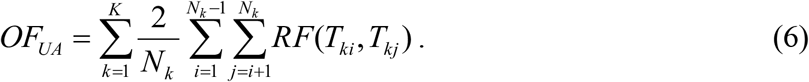

Clearly, the use of the approximation functions defined by Equations (4-6) will not increase the time complexity of our clustering method, which will remain *O*(*nN* ^2^ + *KNI*). The use of these approximation functions should imply the appropriate changes in the formulas of the considered cluster validity indices. For instance, in case of the Caliński-Harabasz index (*CH*) and the objective function *OF*_*MA*_, the intergroup evaluation index *SS*_*B*_ and the intragroup evaluation index *SS*_*W*_ should be calculated as follows:

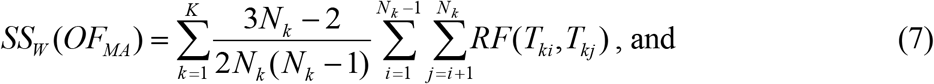

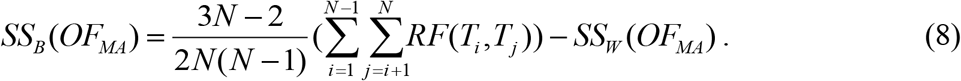

Importantly, the Euclidean objective function *OF*_*EA*_ and the upper bound objective function *OF*_*UA*_ defined in Equations 2 and 6, respectively, differ only by a constant multiplier. This means that they reach their minimum at the same point (i.e. the same tree clustering). Therefore, only one of these functions (i.e. *OF*_*EA*_) was tested in our simulations along with *OF*_*LA*_ and *OF*_*MA*_.

### Clustering trees with different sets of leaves - the supertree approach

In this section, we explain how the tree clustering method introduced above for the case of trees defined on the same set of leaves could be extended to trees whose sets of leaves can differ, as it is often the case in phylogenetic studies, including the famous Tree of Life project (Maddison et al. 2007).

Let Π be a set of *N* unrooted phylogenetic trees that may contain different, but mutually overlapping, sets of labeled leaves. In this case, the original objective function *OF* (Equation 1) can be reformulated as follows:

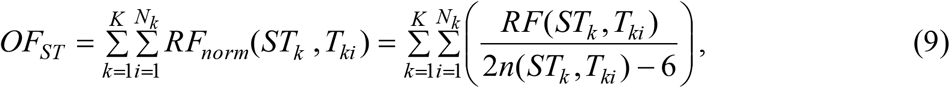

where *K* is the number of clusters, *N*_*k*_ is the number of trees in cluster *k, RF*_*norm*_ (*ST*_*k*_, *T*_*ki*_) is the normalized Robinson and Foulds topological distance between tree *i* of cluster *k*, denoted *T*_*ki*_, and the majority-rule supertree of this cluster, denoted *ST*_*k*_, reduced to a subtree having all leaves in common with *T*_*ki*_. The reduced version of the supertree *ST*_*k*_ is obtained after removing from it all leaves that do not belong to *T*_*ki*_and collapsing the corresponding branches. The *RF* distance is normalized by dividing it by its maximum possible value, which is 2*n*(*ST*_*k*_, *T*_*ki*_) − 6, where *n*(*ST*_*k*_, *T*_*ki*_) is the number of common leaves in trees *ST*_*k*_ and *T*_*ki*_. The normalization is carried out to account equally the contribution of each tree to clustering. Obviously, Equation (9) can be considered only if the number of common leaves in *ST*_*k*_ and *T*_*ki*_ is greater than 3.

We propose to use the following analogue of the Euclidean approximation function (see Equation 2) to avoid supertree computations at each step of *k*-means:

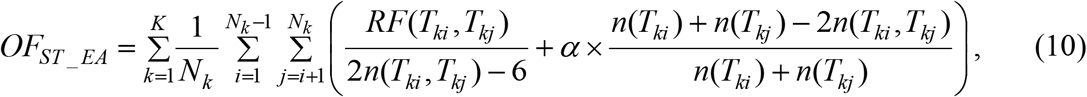

Where *n*(*T*_*ki*_) is the number of leaves in tree *T*_*ki*_, *n*(*T*_*kj*_) is the number of leaves in tree *T*_*kj*_, *n*(*T*_*ki*_, *T*_*kj*_) is the number of common leaves in trees *T*_*ki*_ and *T*_*kj*_, and *α* is the penalization (tuning) parameter, taking values between 0 and 1, used to prevent from putting to the same cluster trees having small percentages of leaves in common. This penalization parameter is necessary in order to get well-balanced clusters in which trees have both high topological and species content similarity. Indeed, the normalized *RF* distance between two large trees can be small only because the trees do not have enough taxa in common. However, such trees should not be necessarily assigned to the same cluster. Equation (10) also implies that two trees belonging to the same cluster have at least four taxa in common, but a higher taxa-similarity threshold can be used as the method’s parameter in order to increase the cluster homogeneity. The objective functions reported in Equations (4-6) and the corresponding cluster validity indices should be normalized in a similar way.

In case of supertree clustering, the *SS*_*W*_ index (see Equation 18) can be computed as follows:

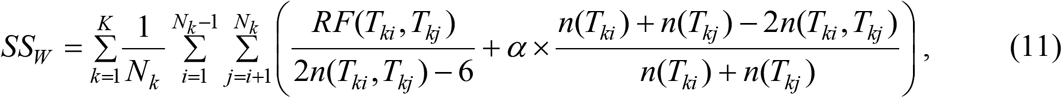

and the *SS*_*B*_ index (see Equation 20) can be computed as follows:

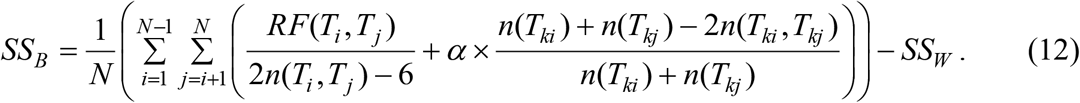

The clustering procedure based on the use of Equation (10) should be carried out with different random input partitions. The best clustering solution can be selected using the value of the adapted *CH* validity index based on Equations (11-12). Once the best clusters of trees are chosen, any existing supertree reconstruction method (Bininda-Emonds 2004) can be applied to infer the related majority-rule supertrees. It is worth noting that Bansal et al. (2010) described a method for building *RF*-based supertrees aiming at minimizing the total *RF* distance between the supertree and the set of input trees. However, they did not normalize the individual *RF* distances. Neither did they consider the possibility of inferring multiple supertrees.

### Program

The program (written in C++) that implements the described method intended for clustering trees and inferring multiple consensus trees and supertrees is freely available at: https://github.com/TahiriNadia/KMeansSuperTreeClustering.

## Results

In this section, we first present our simulation design and discuss the results of the simulation study conducted with synthetic data. Later on we describe the results of our experiments with the SARS-CoV-2 data, originally examined by Lam et al. (2020) and Makarenkov et al. (2021).

### Simulation design

We tested our new method for computing multiple consensus trees and supertrees using the following simulation protocol that included three different simulation experiments. The first experiment involved partitioning phylogenies having identical sets of leaves (i.e. multiple consensus trees were constructed) and included a comparison of our new method with some state-of-the-art tree partitioning algorithms. The second experiment consisted in partitioning phylogenies having different sets of leaves (i.e. multiple consensus supertrees were constructed) without using penalization in the objective function (i.e. the value of the penalization parameter *α* in Equation 10 was set to 0). The third experiment involved partitioning phylogenies having different sets of leaves using the penalization parameter *α* (its value varied between 0 and 1) in the objective function (29). The detailed simulation protocol adopted in our study is presented in Appendix A4.

### Simulation results

The results of our simulations conducted with synthetic data are illustrated in Figures 1 and 2 (clustering of gene trees defined on the same set of taxa), and Figures A3, A4 (Appendix A.5) and 3 (clustering of gene trees defined on different, by mutually overlapping, sets of taxa). Figure 1 presents the clustering performances provided by the six compared variants of our partitioning algorithm (see Appendix A.4 and the legend of Fig. 1). The best overall results in this simulation were provided by the variant of our algorithm based on the *OF*_*EA*_ objective function using the approximation by Euclidean distance and *CH* cluster validity index (*CH-E*). The results provided by the variant based on the *OF*_*MA*_ objective function with approximation by the mean of interval and *CH* cluster validity index (*CH-MI*) were only slightly worse. At the same time, the average ARIs (Adjusted Rand Indices) obtained for the variants based of the *BH* and *Gap* statistics were generally much lower than those provided by the variants based on the *CH* and Silhouette cluster validity indices. Among the algorithm’s variants that were able to deal with homogeneous data (when the number of clusters *K* was equal to 1), the variant based on *BH* outperformed that based on *Gap* for the lower numbers of clusters (*K* = 1, 2 and 3), but was less effective than it for the higher numbers of clusters (*K* = 4 and 5). It is worth noting that the results provided by the variants based on the *CH* index (*CH-E, CH-MI* and *CH-LB*) are very stable; they do not vary a lot depending on the number of clusters, the number of leaves and the coalescence rate.

**Fig. 1.**
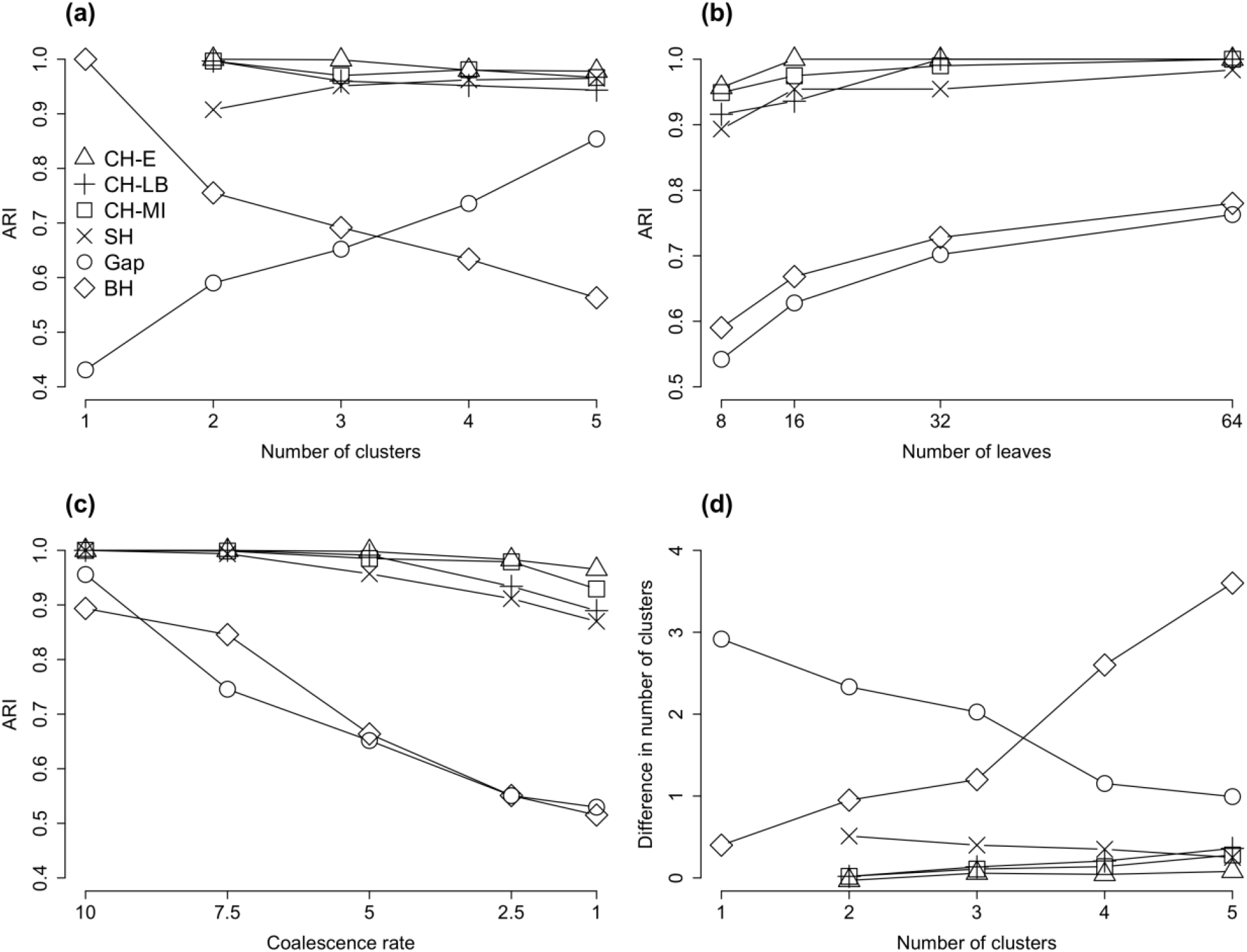
Classification performance of the six variants of our *k*-means tree clustering algorithm applied to trees with identical sets of leaves using, respectively: *OF*_*EA*_ objective function with approximation by Euclidean distance and *CH* cluster validity index (*CH-E*), *OF*_*LA*_ objective function with approximation by the lower bound and *CH* cluster validity index (*CH-LB*), *OF*_*MA*_ objective function with approximation by the mean of interval and *CH* cluster validity index (*CH-MI*), *OF*_*EA*_ objective function and Silhouette cluster validity index (*SH*), *OF*_*EA*_ objective function and Gap cluster validity index (*Gap*), and *OF*_*EA*_ objective function and Ball and Hall cluster validity index (*BH*). Only the *Gap* and *BH* indices could be used to assess the algorithm’s performance on datasets containing one cluster. The results are presented in terms of average *ARI* with respect to the: (a) number of tree clusters, (b) number of tree leaves and (c) coalescence rate, and in terms of the: (d) average absolute difference between the true and the obtained number of clusters. The coalescence rate parameter in the HybridSim program was set to 5 in simulations (a), (b) and (d). The presented results are the averages taken over all considered numbers of leaves in simulations (a), (c) and (d), and all considered numbers of clusters in simulations (b) and (c).

**Fig. 2.**
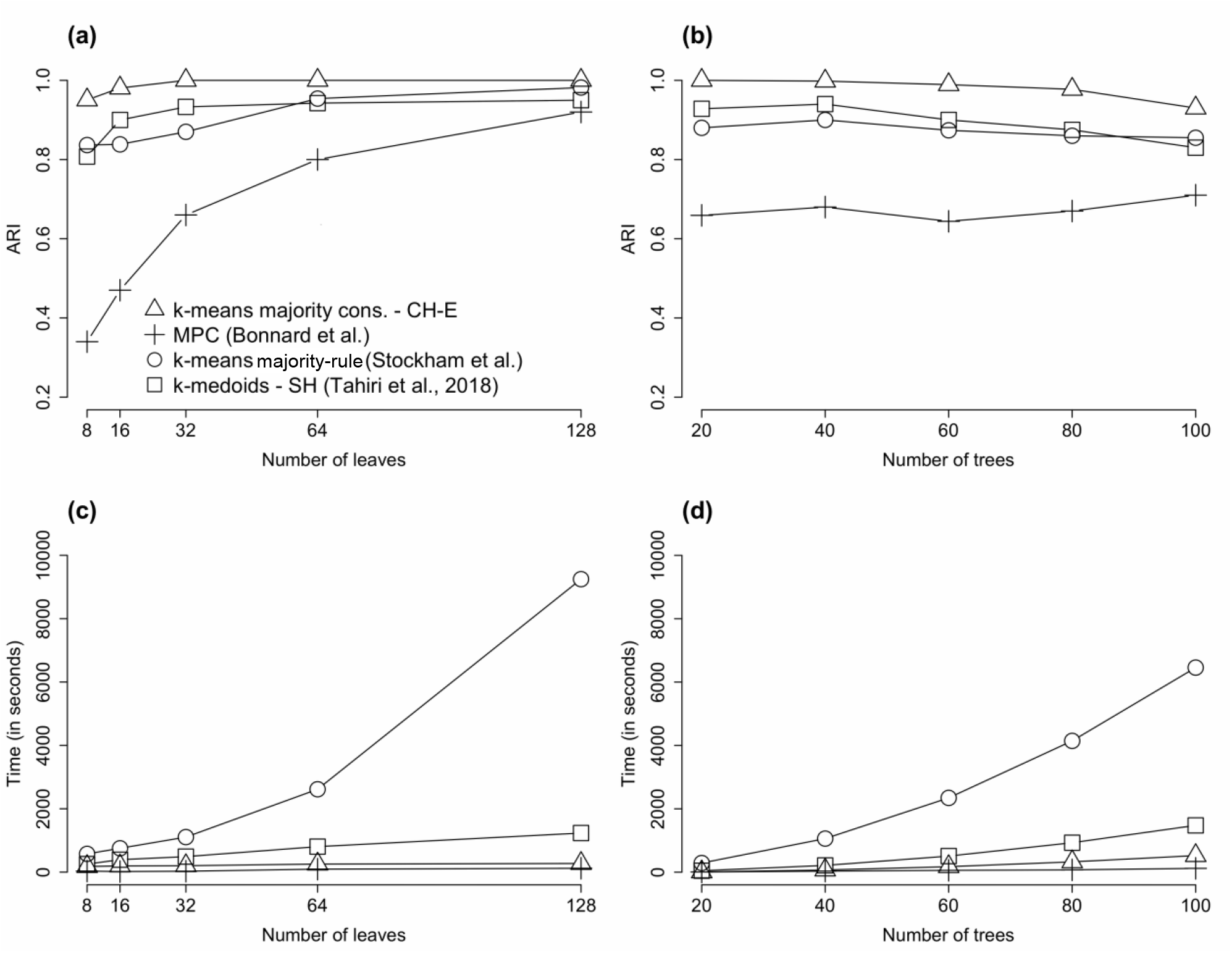
Comparison of our algorithm (Δ) based on *k*-means tree clustering with *OF*_*EA*_ objective function and *CH* cluster validity index (*CH-E*), the MPC tree clustering algorithm (+) by Bonnard et al. (2006), the tree clustering algorithm by Stockham et al. (2002) (○) based on *k*-means clustering with squared *RF* distance, and the *k*-medoids tree clustering algorithm by Tahiri et al. (2018) based on the *RF* distance and *SH* cluster validity index (□). The comparison was made in terms of the average *ARI* (panels a and b) and the average running time (measured in seconds) of the algorithms (panels c and d) with respect to the number of leaves and trees by cluster. The coalescence rate parameter was set to 5 and the number of clusters varied from 2 to 5 in this simulation.

Figure 2 presents the results of comparison of our most stable algorithm’s variant, *CH-E*, with the state-of-the-art tree clustering methods, including the MPC tree clustering algorithm by Bonnard et al. (2006), the tree clustering algorithm by Stockham et al. (2002) that is based on the squared *RF* distance and recomputing the majority-rule consensus trees at each iteration of *k*-means, and the *k*-medoids tree clustering algorithm by Tahiri et al. (2018) based on the *RF* distance (non-squared) and the *SH* cluster validity index. The curves presented in Figure 2 (a and b) indicate that our *CH-E* strategy clearly outperformed the three other competing methods in terms of the clustering quality (i.e. average ARI results). The clustering results provided by the methods of Stockham et al. and Tahiri et al. were very close, and both of them generally outperformed the MPC approach. However, in case of large trees (with 128 leaves) the average ARI results provided by the four tested methods were close. All the methods, but especially MPC, show an increase in the ARI values as the number of tree leaves grows. Moreover, the *CH-E* and MPC algorithms were by far the best methods in terms of the running time for both simulation parameters considered: the number of tree leaves (Fig 2c) and the number of trees (Fig 2d). These results suggest that our new algorithm, along with MPC, is well suited for the analysis of large phylogenetic datasets. However, for smaller phylogenies (with < 128 leaves) our *CH-E* strategy represents the best choice overall.

Classification performances of our supertree clustering algorithm based on the *OF*_*ST_EA*_ objective function used with *CH* and *BH* cluster validity indices are presented in Appendix A4 (see Figs. A3 and A4, respectively).

Figure 3 illustrates the performance of our supertree clustering algorithm applied to gene trees with different numbers and sets of leaves using the *OF*_*ST_EA*_ objective function with approximation by Euclidean distance and *CH* cluster validity index. In this experiment, the value of the penalization parameter *α* in Equation 10 varied from 0 to 1, with the step of 0.2. Obviously, when no species are removed from the dataset, the penalization term in Equation 10 equals 0 and has no impact on the clustering performance. However, in all other cases, the presented ARI curves showed different behaviour and reached its maxima at different values of *α* (i.e. for the case of 10% of missing species the maximum value of ARI was reached at *α* = 0, for the case of 25% of missing species the maximum was reached at *α* = 0.2, with very close ARI results obtained for *α* = 0.4 and 1.0, whereas for the case of 50% of missing species the maximum was reached at *α* = 1.0). Thus, we can conclude that the value of the penalization parameter *α* must be chosen with respect to the number of missing taxa in the input trees.

**Fig. 3.**
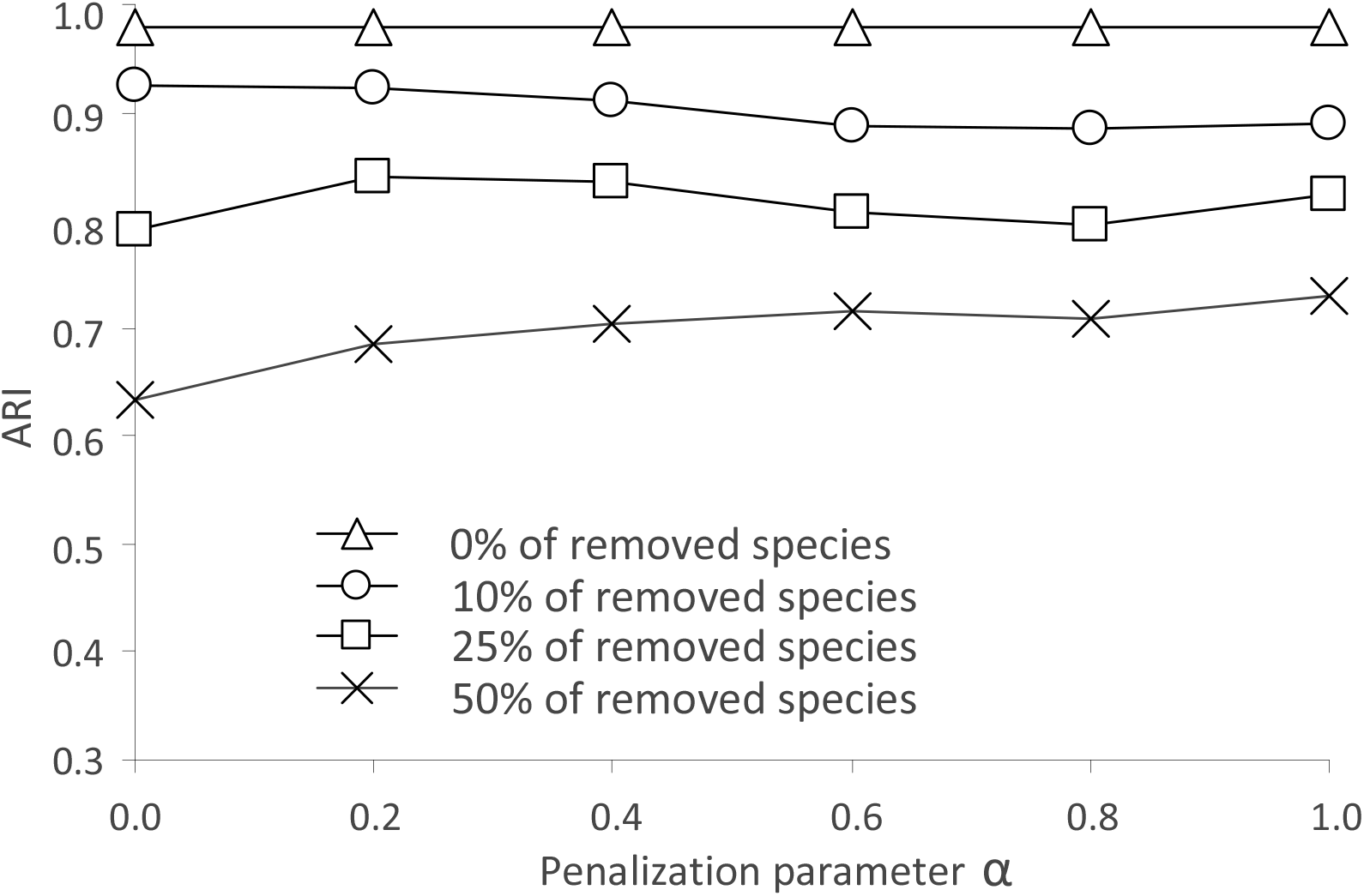
Classification performance of our *k*-means-based supertree clustering algorithm applied to trees with different numbers and sets of leaves using the *OF*_*ST_EA*_ objective function with approximation by Euclidean distance and *CH* cluster validity index. The results are presented in terms of *ARI* with respect to the value of the penalization parameter *α*, which varied between 0 and 1 with the step of 0.2. The coalescence rate parameter in the HybridSim program was set to 5 in this simulation. The presented results are the average ARIs computed over all numbers of leaves and clusters considered in our simulations.

## Exploring the patterns of evolution of SARS-CoV-2 genes

In this section, we first describe the SARS-CoV-2 dataset examined in our work, then give some details regarding the applied multiple sequence alignment and tree inference methods, and finally present and discuss the results of our supertree clustering analysis.

### Data description

To carry out our the supertree clustering analysis, we considered the case of evolution of 43 betacoronavirus organisms, including : (1) four SARS-CoV-2 genomes from China, Australia, Italy and USA, coming from different clades of the Gisaid SARS-CoV-2 phylogeny (see https://www.gisaid.org; Shu and McCauley 2017); (2) the RaTG13 bat CoV genome from *Rhinolophus affinis* (data collected in the Yunnan province of China); (3) five Guangxi (GX) pangolin CoV genomes (data provided by the Beijing Institute of Microbiology and Epidemiology); (4) two Guangdong (GD) pangolin CoV genomes (data extracted from dead Malayan pangolins in the Guangdong province of China); (5) the bat CoV ZC45 and ZXC21 genomes (i.e. bat CoVZ clade with data coming from the Zhejiang province of China); (6) five additional CoV genomes extracted from bats across different provinces of China (i.e. BtCoV 273 2005, Rf1, HKU3-12, HKU3-6 and BtCoV 279 2005 CoVs); (7) four SARS-CoV strains related to the first SARS outbreak (i.e. SARS, Tor2, SARS-CoV BJ182-4 and bat Rs3367 CoV found in *Rhinolophus sinicus*); (8) the BtKY72 and BM48 31 BGR 2008 CoV genomes, extracted from bats in Kenya and Bulgaria; (9) the MERS-CoV and the related bat HKU-4 and HKU-5 CoV genomes; (10) a human HKU1 CoV genome; (11) a feline CoV genome; (12) four murine CoV and the related Rat Parker CoV genomes; (13) an equine CoV genome; (14) a porcine CoV genome; (15) the rabbit HKU14 CoV genome; (16) human enteric and human OC43 CoV genomes; (17) three bovine CoV genomes, including AH187 and OH440 bovine CoVs. The first 25 of these CoV genomes (sub-groups 1 to 8 above) include the closest relatives of SARS-CoV-2; they have been originally examined in Lam et al. (2020). The remaining 18 CoVs (sub-groups 9 to 17 above) comprise betacoronaviruses labeled as common cold CoVs in the Gisaid coronavirus tree (Shu and McCauley 2017) and those studied by Prabakaran et al. (2006). Supplementary Table 1 (see Supplementary Material) provides the organism names, host species, and Gisaid or GenBank accession numbers for the 43 coronaviruses considered in our study.

### Methods details

Our analysis was conducted for 11 main genes of the SARS-Cov-2 genome (i.e. genes ORF1ab, S, ORF3a, E, M, ORF6, ORF7a, ORF7b, ORF8, N and ORF10) as well as for the RD domain of the spike protein because of its key evolutionary importance. Thus, 12 gene phylogenies were inferred and analyzed in our study. It is worth mentioning that some gene annotations were absent in the GenBank and Gisaid databases (i.e. annotations for genes ORF6, ORF7a, ORF7b, ORF8 and ORF10 for taxa from sub-groups 9 to 17 and annotations for gene ORF8 for taxa from sub-group 8 were unavailable), thus leading to gene phylogenies having different, but mutually overlapping, sets of leaves.

The VGAS program (Zhang et al. 2019), intended to identify viral genes and carry out gene function annotation, was executed to validate all betacoronavirus genes found in GenBank and Gisaid. We then performed multiple sequence alignments (MSAs) for the 11 considered coronavirus genes (DNA sequences) and for the RB domain (amino acid sequences) by means of the MUSCLE algorithm (Edgar 2004) using the default parameters of the MegaX program (v. 10.1.7) (Kumar et al. 2018). The obtained MSAs were used to build gene trees presented in Supplementary Figures 1 to 12 (see Supplementary Material). Moreover, in order to apply the HGT and recombination detection methods on the obtained gene and consensus trees, we inferred a species phylogeny (see Fig. 4d) of the 43 considered CoVs using the same version of MUSCLE. The GBlocks algorithm (v. 0.91b, Castresana 2000), available at the Phyloge-ny.fr web server (Dereeper et al. 2008), was then used with the less stringent correction option to remove MSA sites with large gap ratios. Gene and genome trees that will be further used in clustering and HGT (Horizontal Gene Transfer) and recombination analyses were inferred using the RAxML algorithm (v. 0.9.0; Stamatakis 2006). The most suitable DNA/amino acid substitution model determined by MegaX, and available on the RAxML web site (https://raxml-ng.vital-it.ch), was used for each MSA. Precisely, the (GTR+G+I) model was found to be the best-fit substitution model for genes ORF1ab, S, N and for the whole genomes, the (HKY+G) model was the most suitable for genes ORF3a, E, ORF6, ORF7a, the (HKY+I) model for gene ORF7b, the (HKY+G+I) for gene ORF8, the (JC) model for gene ORF10, and the (WAG+G) model for the RB domain. The tree inference was conducted using the bootstrap option (with 100 replicates for each MSA considered).

**Fig. 4.**
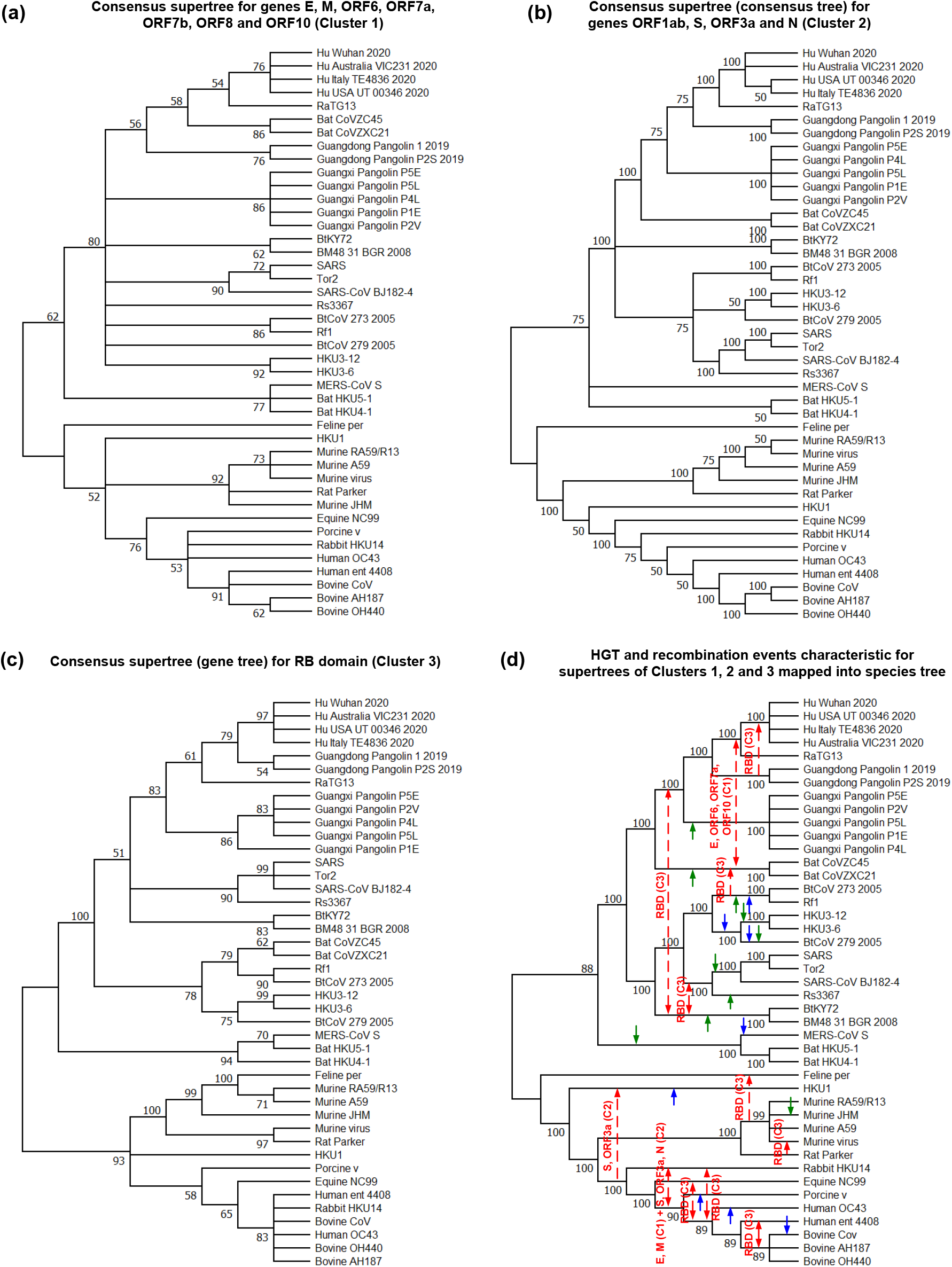
Three consensus supertrees illustrating three main ways of evolution of betacoronavirus genes and HGT-recombination network depicting the most important gene transfer and recombination events found for each consensus supertree. (a) Supertree of Cluster 1 is the best heuristic search (*hs*) CLANN supertree inferred for the phylogenies of genes E, M, ORF6, ORF7a, ORF7b, ORF8 and ORF10; (b) Supertree of Cluster 2 is the extended majority-rule consensus tree inferred for the phylogenies of genes ORF1ab, S, ORF3a and N (defined on the same set of 43 taxa); (c) Supertree of Cluster 3 is the RAxML gene tree of the RB domain of the spike protein (a unique member of this cluster); (d) Horizontal gene transfer and recombination events inferred by the HGT-Detection program for each of the three consensus supertrees. Bootstrap scores are indicated on the internal tree branches. Branches with bootstrap support lower than 50% were collapsed. Transfer directions are represented by arrows (when the direction is uncertain, the arrow is bidirectional). Gene(s) affected by the transfer and the corresponding tree cluster number are indicated on the red arrows. Blue (for Cluster 1) and green (for Cluster 2) arrows represent additional HGT events found by the HGT-Detection program for individual genes of these clusters (these events cannot be inferred directly from consensus supertrees).

The consensus supertrees (see Fig. 4a, b and c) for each cluster found by our algorithm were inferred using the CLANN program (Creeve and McInerney 2005). The HGT-Detection program (Boc et al. 2010) from the T-Rex web server (Boc et al. 2012) was used to infer directional horizontal gene transfer-recombination network (Huson and Bryant 2005; Beiko et al. 2005) for the three obtained consensus supertrees (see Fig. 4d).

Multiple sequence alignments for all gene and genome sequences used in this study as well as all inferred gene and species phylogenies (in the Newick format) are available at: http://www.info2.uqam.ca/~makarenkov_v/Supplementary_Material_data.zip.

### Supertree clustering and HGT-recombination analysis of SARS-CoV-2 data

We performed the CoV gene tree clustering to identify genes having similar evolutionary patterns. The analysis was conducted using the supertree clustering algorithm described in the Materials and Methods section. In total, 7 gene trees with 43 leaves (the phylogenies of genes ORF1ab, S, ORF3a, E, M, N and that of the RB domain), 4 gene trees with 25 leaves (the phylogenies of genes ORF6, ORF7a, ORF7b and ORF10) and 1 gene tree with 23 leaves (the phylogeny of gene ORF8) were considered. The internal braches of gene phylogenies with bootstrap support lower than 50% were collapsed prior to conducting gene tree clustering. The average number of missing leaves by gene tree was equal to 17.8% According to our simulations (see Fig. 3), the optimal value of the penalization parameter *α* could vary between 0 and 0.2 for such data. Here, we present the clustering results obtained with the value of *α* = 0.1 (very similar clusterings were obtained with *α* = 0 and *α* = 0.2; only one tree changes its cluster membership with *α* = 0.2). The obtained clustering solution with 3 disjoint clusters is presented in Figure 4 (a, b and c). This solution encompasses three main patterns of evolution of betacoronavirus genes. The phylogenies of genes E, M, ORF6, ORF 7a, ORF7b, ORF8 and ORF10 were assigned to Cluster 1, those of genes ORF1ab, S, ORF3a and N to Cluster 2, and that of the RB domain to Cluster 3. The consensus supertrees for each cluster were then inferred using the CLANN program (Creeve and McInerney 2005). The supertree for Cluster 1 (Fig. 4a) was inferred using the heuristic search (*hs*) and *bootstrap* (performed with 100 replicates) options available in CLANN. The supertree for Cluster 2 (Fig. 4b) was obtained using the *consensus* option of CLANN as all four trees forming this cluster contains 43 taxa. The consensus supertree of Cluster 3, containing a singleton element (i.e. the gene phylogeny of the RB domain) was its RAxML gene tree (Fig. 4c).

Moreover, we also conducted a detailed HGT and recombination analysis of the obtained tree cluster supertrees in order to identify the main HGT and recombination patterns characterizing the evolution of SARS-CoV-2 and related betacoronaviruses. HGT and recombination are widespread reticulate evolutionary processes contributing to diversity of betacoronaviruses, as well as of most other viruses, allowing them to overcome selective pressure and adapt to new environments (Pérez-Losada et al. 2015). The HGT-Detection algorithm of Boc et al. (2010) was carried out independently for each of the three consensus supertrees inferred for our data. The obtained gene transfers, representing common HGT and recombination trends of the tree cluster under study, are indicated by red arrows in Figure 4d. We then completed our analysis by conducting an independent gene transfer detection for all individual gene trees included in Clusters 1 and 2 (the detected individual gene HGTs that were different from the previously recovered common transfers are indicated by blue and green arrows, respectively).

The obtained clustering and HGT detection results highlight the uniqueness of the evolution of the RB domain. They also suggest that the RB domain of SARS-CoV-2 could be acquired by a horizontal transfer of genetic material, followed by intragenic recombination, from the Guangdong pangolin CoV. Furthermore, they emphasize the stability of evolutionary patterns of the longest CoV genes (i.e. ORF1ab, S, ORF3a and N assigned to Cluster 2) as the consensus supertree (i.e. consensus tree in this case) of this cluster has a well-resolved structure.

The consensus supertree of Cluster 1 is less resolved of all supertrees what could be expected since this cluster contains 7 of 12 gene trees considered in our study. The topology of the Cluster 1 supertree points out that SARS-CoV-2 could not only be a mosaic created by recombination of the bat RaTG13 and Guangdong pangolin CoVs, but is also a very close relative of the bat ZC45 and ZXC21 CoV strains, which are the most closely located taxa to the clade of the RaTG13 and SARS-CoV-2 viruses in this supertree.

## DISCUSSION

### Main properties and advantages of the presented method

Consensus tree and supertree inference methods synthesize collections of gene phylogenies into comprehensive trees that preserve main topological features of the input phylogenies and include all taxa present in them. In this paper, we introduced a new systematic method for inferring multiple alternative consensus trees and supertrees from a given set of phylogenetic trees, which can be defined either on the same set of taxa (case of multiple consensus trees) or on different sets of taxa with incomplete taxon overlap (case of multiple supertrees). To the best of our knowledge, the problem of building multiple alternative supertrees has not been addressed yet in the literature. The inferred alternative consensus trees and supertrees represent the most important evolutionary patterns characterizing the evolution of genes under study. They are generally much better resolved than a single consensus tree or a single super-tree inferred by traditional methods. Thus, a multiple consensus tree or supertree inference approach has the potential to build supertrees that retain much more plausible information from the input set of gene phylogenies. A single consensus tree or supertree could be an appropriate representation of a given set of gene trees only if all of them, or a large majority of them, follow the same evolutionary patterns. For example, the presented method allows one to identify ensembles of genes that underwent similar horizontal gene transfer, hybridization or intragenic/intergenic recombination events, or those that were affected by similar ancient duplication events during their evolution. As we showed in the Results section, our method can be effectively used to retrace main evolutionary patterns of SARS-CoV-2 genes. It could be used as well for inferring alternative subtrees of Tree of Life.

The presented method relies on multiple runs of the *k*-means partitioning algorithm applied to the non-squared Robinson and Foulds distances (original or normalized) between the input trees. A number of efficient approximations of the straightforward objective function (Equation 1), preventing us from computing a consensus tree for any considered cluster of trees in the internal loop of *k*-means and using some remarkable properties of the *RF* distance, have been introduced (see Equations 2 and 4-6). These equations allows us to precompute all *RF* distances prior to carrying out the *k*-means tree clustering and ensure that a basic *k*-means object relocation operation, consisting in removing a given tree *T* from its current cluster *C* and assigning it to the best possible cluster different from *C* (if any), can be performed in *O*(*K*), where *K* is the number of tree clusters. This property makes our method perfectly suitable for analysis of large evolutionary datasets. In order to compute the *RF* distance between pairs of trees defined on different, but mutually overlapping, sets of leaves we propose to reduce them pairwise to common sets of leaves and then to normalize the obtained distance value. In case of supertree clustering, we also added to the objective function of the method the term including the penalization parameter *α*, which is used to create well-balanced clusters that contain trees with both high topological and species content similarity. The use of the *RF* distance, and not of its quadratic form, in tree clustering is justified by the fact that the majority-rule consensus tree of a set of trees is a median tree of this set in the sense of the *RF* distance (Barthélemy and McMorris 1986). Moreover, Bansal et al. (2010) showed that *RF*-based supertrees are supertrees that are consistent with the largest number of clades from the input trees. We also showed how the popular Caliński-Harabasz (*CH*), Silhouette (*SH*), Ball and Hall (*BH*), and Gap cluster validity indices could be adapted to tree clustering with *k*-means. The *CH* and *SH* indices are suitable for clustering heterogeneous data (when the number of clusters *K* ≥ 2), whereas the *BH* and Gap indices can be used to cluster both homogenous (when *K* = 1) and heterogeneous data. Using simulations, we demonstrated that the version of our method based on Euclidean approximation (Equation 2) typically outperforms the existing methods for building multiple alternative consensus trees, such as MPC (Bonnard et al. 2006), *k*-means tree clustering with squared *RF* distance by Stockham et al. (2002) and *k*-medoids tree clustering by Tahiri et al. (2018), in terms of both clustering quality and running time.

### Future extensions of the method

The statistical robustness of phylogenetic trees is a very important factor that should not be neglected when inferring and interpreting the results of phylogenetic analysis. We know that the *RF* distance is twice the number of bipartitions present in one of tree and absent in the other (Robinson and Foulds 1981). Thus, we can incorporate bootstrap scores of the input trees in the computation by considering the following objective function in the framework of consensus tree clustering:

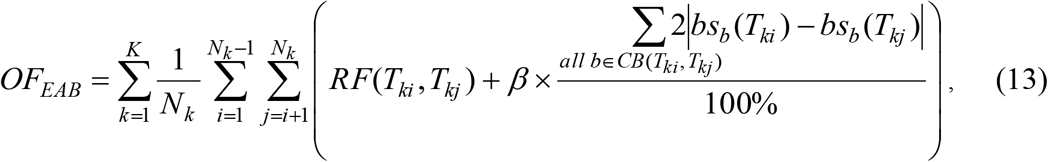

Where *bs*_*b*_ (*T*_*ki*_) and *bs*_*b*_ (*T*_*kj*_) are the bootstrap scores, expressed in percentages, of internal branch *b* that induces a common bipartition in trees *T*_*ki*_ and *T*_*kj*_, *CB*(*T*_*ki*_, *T*_*kj*_) is the set of all common internal branches (i.e. branches inducing the same bipartitions) in *T*_*ki*_ and *T*_*kj*_, and *β*is the penalization parameter, taking values between 0 and 1, used to penalize pairs of input trees with different bootstrap scores of their internal branches inducing common bipartitions. Based on Equation 13, trees with similar bootstrap support of their internal branches inducing the same bipartitions of taxa will have a greater potential to be assigned to the same cluster. It is worth mentioning that branch lengths of compared branches can be taken into account in a similar way in the objective function of the clustering algorithm. In this case, the absolute difference between bootstrap scores of the two compared branches inducing the same bipartition in *T*_*ki*_ and *T*_*kj*_ (see Equation 13) can be replaced by the absolute difference between their lengths, whereas the maximum of the two branch lengths will replace 100% in the fraction denominator.

We can also use the following analogue of the Euclidean approximation function (see Equation 10) to take into account bootstrap support of the internal tree branches and avoid super-tree computations at each step of the supertree *k*-means clustering:

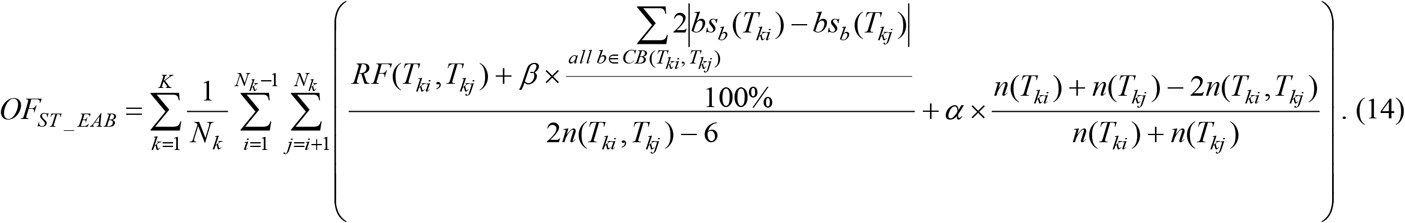

Another interesting option for further investigations concerns the use of other popular tree distances in the objective function of the clustering algorithm. In this context, the two most promising of them seem to be the *Branch score distance* (Kuhner and Felsenstein 1994) and the *Quartet distance* (Bryant et al. 2000).

The branch score distance is defined as follows. Let us consider two phylogenetic trees *T* and *T’* defined on the same set of *n* taxa and the large set (*BP*_1_, *BP*_2_, …, *BP*_*NB*_) of all possible bipartitions existing for these taxa. For each tree, we can determine a large vector of non-negative values **bp** = (*b*_1_, *b*_2_, …, *b*_*NP*_), in which *b*_*i*_ is equal to the branch length of the branch corresponding to bipartition *BP*_*i*_ if this bipartition exists in the tree, otherwise it is equal to 0. For two trees *T* and *T’*, whose bipartition vectors are **bp** and **bp**’, the branch score distance is defined as the squared Euclidean distance between these vectors: 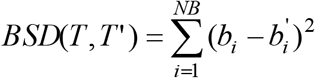, where *NB* is the number of all possible existing bipartitions.

The quartet distance (*QD*) is defined as the number of quartets of tree leaves that induce a subtree topology that occur in only one of the two compared trees. According to its definition, the quartet distance is a symmetric difference distance. Thus, its square root (*QD*^1*/*2^) is Euclidean (Critchley and Fichet 1994) and an analogue of Equation 2, in which *RF* is replaced by *QD*, can be used in the objective function of the clustering algorithm. Another advantage of the quartet distance is that it can be computed in *O*(*n*log*n*) (Brodal et al. 2004), where *n* is the number of leaves in both trees involved in computation.

Another tree distance which could be suitable for tree clustering is the *Billera–Holmes– Vogtmann (BHV) distance* (Billera et al. 2001). The *BHV* distance between weighted trees is defined as the geodesic, or shortest path, distance inside treespace in which trees are viewed as (2*n*−3)-dimensional vectors of their bipartition weights within the larger (2^*n*−1^−1)-dimensional space of all graphs (St. John 2017). The *BHV* distance between two trees can be computed in *O*(*n*^4^) (Owen and Provan 2010) and approximated in linear time (Amenta et a l. 2007). The continuous treespace introduced by Billera et al. (2001) provides a perfect environment for computation of average trees, while the classical Euclidean mean, when applied to tree vectors, can yield vectors not corresponding to trees (St. John 2017). In the *BHV* framework, the majority consensus tree (McMorris et al. 1983) corresponds to the mode and the Fréchet mean corresponds to the average of a given set of trees. The Fréchet mean tree of a given set of *N* trees is the tree that minimizes the sum of the squared *BHV* distances to the given set Π = {*T*_1_, *T*_2_, …, *T*_*N*_} of *N* phylogenetic trees defined on the same set of taxa, i.e. 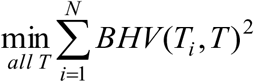. A normalized sum of such minimum values computed independently for each considered cluster of trees can constitute an objective function of a tree clustering method based on the *BHV* distance. Fortunately, the Fréchet mean tree is unique and its approximation can be calculated in polynomial time by an iterative algorithm (Sturm 2003). However, the Fréchet mean may also demonstrate a non-Euclidean sticky behaviour, as changing an input tree does not necessarily change the mean tree of the dataset, in contrast to Euclidean space (Miller et al. 2015 ; St. John 2017). Alternatively, the median (which is a more robust estimator than the mean) of a set of trees can also be considered when clustering tree. The *BHV* distance median of a set of trees is the tree that minimizes the sum of non-squared *BHV* distances to those trees (see Benner et al. 2014 for an algorithm computing the *BHV* median).

Another widely-used criterion for assessing differences between trees is the *Subtree Prune-and-Regraft* (*SPR*) distance (Hein et al. 1996). This distance is integrated in a variety of tree building methods exploring different tree topologies (Gascuel 2005). Moreover, Whidden et al. (2014) determined that the *SPR* distance can be used to build supertrees and that *SPR*-based supertrees are significantly more similar to the known species history than *RF*-based supertrees given biologically plausible rates of simulated horizontal gene transfers. The problem of computing the *SPR* distance between two trees is NP-hard (Bordewich and Semple 2005). However, in practice, its approximation can be computed using a fixed-parameter-bounded search tree algorithm in combination with a linear-time formulation of Linz and Semple’s cluster reduction to solve an equivalent maximum agreement forest problem (Linz and Semple 2011; Whidden et al. 2014). Nevertheless, the *SPR* distance, as well as the other popular tree topology rearrangement distances such as *Nearest Neighbor Interchange* (*NNI*) and *Tree Bisection and Reconnection* (*TBR*), has no Euclidean properties and new formulas and algorithms should be designed in order to adapt it to tree clustering.

## SUPPLEMENTARY MATERIAL

Supplementary material is available for this manuscript.

## ACKNOWLEDGMENTS

We would like to thank Nicolas Lartillot for providing us with the MPC tree clustering program, Pierre Legendre and Bogdan Mazoure for helping us with the analysis of SARS-CoV-2 data, and Louise Laforest and Alexander Friedmann for insightful conversations and helpful comments. We also thank Compute Canada for providing access to high-performance computing facilities.

## Appendix A

### A.1 *RF* is not a Euclidean distance, but its square root is Euclidean

The following counter-example, involving four trees *T*_1_, *T*_2_, *T*_3_ and *T*_4_ with five leaves each (see Fig. A1), can be used to show that *RF* is not Euclidean. It is the simplest example possible because for the trees with four leaves, the *RF* distance has the Euclidean properties.

**Fig. A1.**
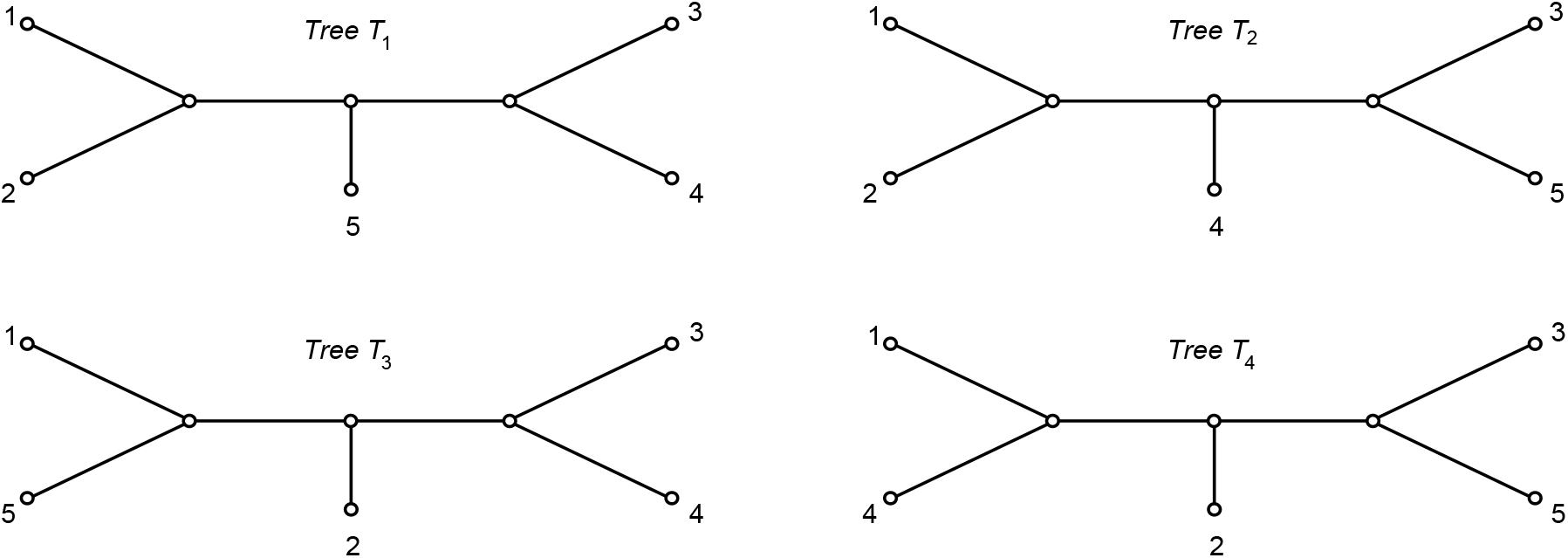
Four unrooted phylogenetic trees, *T*_1_, *T*_2_, *T*_3_ and *T*_4_ with five leaves used to show that the Robinson and Foulds topological distance is not a Euclidean distance.

The 1-bipartitions, corresponding to tree leaves, are common to the four trees in Figure A1. The 2-bipartitions here, defined by the subsets of two taxa, are as follows: *T*_1_: {1, 2} and {3, 4}, *T*_2_: {1, 2} and {3, 5}, *T*_3_: {1, 5} and {3, 4}, and *T*_4_: {1, 4} and {3, 5}. Therefore, *RF*(*T*_1_, *T*_2_) = 2, *RF*(*T*_2_, *T*_3_) = 4 and *RF*(*T*_1_, *T*_3_) = 2. In a Euclidean case, this would place the tree *T*_1_ in the middle of the interval [*T*_2_,*T*_3_] (see Fig. A2). At the same time, we know that *RF*(*T*_1_,*T*_4_) = 4 and *RF*(*T*_3_,*T*_4_) = 4, meaning that *T*_4_ should be located on the perpendicular bisector of the interval [*T*_1_,*T*_3_]. However, this contradicts the fact that *RF*(*T*_2_,*T*_4_) = 2.

**Fig. A2.**
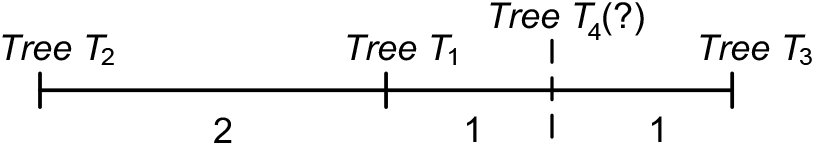
An illustration depicting the position of the four trees from Figure A1 used to show that the Robinson and Foulds distance is not a Euclidean distance.

We will now recall a few mathematical results which will be useful in the sequel. They will allow us to suggest a new clustering strategy via the square root of the *RF* distance (see the main text). In a series of beautiful papers, Kelly and Deza introduced hypermetric spaces and then extended them to quasi-hypermetric spaces (Kelly 1972; Deza and Laurent 1997). Quasi-hypermetric spaces satisfy a so-called inequality of negative type, i.e. given a metric space (*X, d*): for every strictly positive integer *k*, for every real numbers *λ*_1_, …, *λ*_*k*_, and for all variables *x*_1_, …, *x*_*k*_ in *X*, the following inequality holds: Σ_*i*=1,.*k*_ Σ_*j* =1,.*k*_ *λ*_*i*_ *λ*_*j*_ *d*(*x*_*i*_, *x*_*j*_) ≤ 0. Then, it has been observed that this inequality corresponds exactly to the Schoenberg condition for the square root (*d*^1*/*2^) of *d* being of the Euclidean type. Kelly (1972) provided several examples of quasi-hypermetric spaces, such as normed spaces and lattices, and established that a symmetric difference distance is hypermetric. Therefore, its square root (*d*^1*/*2^) is Euclidean (see Critchley and Fichet (1994) for more details). Because the Robinson and Foulds distance is a symmetric difference distance (Robinson and Foulds 1981), its square root (*RF*^1*/*2^) is Euclidean.

### A.2 Caliński-Harabasz, Silhouette, Gap and Ball-Hall cluster validity indices adapted for tree clustering with *k*-means

#### Caliński-Harabasz cluster validity index adapted for tree clustering with k-means

The first cluster validity index we consider here is the Caliński-Harabasz index (Caliński and Harabasz 1974). This index, sometimes called the variance ratio criterion, is defined as follows:

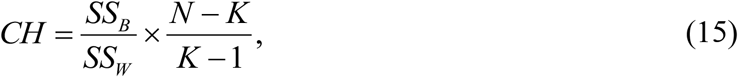

where *SS*_*B*_ is the index of intergroup evaluation, *SS*_*W*_ is the index of intragroup evaluation, *K* is the number of clusters and *N* is the number of objects (i.e. trees in our case). The optimal number of clusters corresponds to the greatest value of *CH*.

In the traditional version of *CH*, when the Euclidean distance is considered, the *SS*_*B*_ coefficient is evaluated by using the *L*^2^-norm:

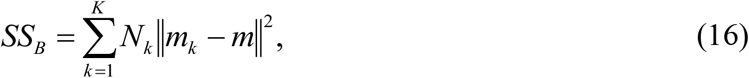

where *m*_*k*_ (*k* = 1 … *K*) is the centroid of cluster *k, m* is the overall mean (i. e. centroid) of all objects in the given dataset *X* and *N*_*k*_ is the number of objects in cluster *k*. In the context of the Euclidean distance, the *SS*_*W*_ index can be calculated using the two following equivalent expressions:

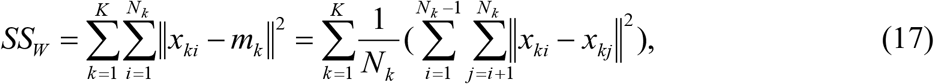

where *x*_*ki*_ and *x*_*kj*_ are the objects *i* and *j* of cluster *k*, respectively (Caliński and Harabasz 1974).

To use the analogues of Equations 16 and 17 in tree clustering, we need to define the concept of centroid for a given set of trees. The *median tree* (Barthélemy and Monjardet 1981; Barthélemy and McMorris 1986) plays the role of this centroid in our tree clustering algorithm. The median procedure is defined as follows (Barthélemy and Monjardet 1981). The median trees, Md(Π), for a given set of trees Π = {*T*_1_, …, *T*_*N*_} having the same set of leaves *S*, is the set of all trees *T* defined on *S*, such that: 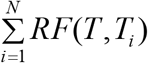 is minimized. If *N* is odd, then the *majority-rule consensus tree*, Maj(Π) of Π, is the only element of Md(Π). If *N* is even, then Md(Π) is composed of Maj(Π) and of some more resolved trees (see Barthélemy and Monjardet 1981 for more details).

We propose to use approximation formulas based on the properties of the Euclidean distance in order to define *SS*_*B*_ and *SS*_*W*_ in *k*-means-like tree clustering. These formulas do not require the computation of the majority (or the extended majority)-rule consensus trees at each iteration of *k*-means. Precisely, we replace the term ‖*x*_*ki*_ − *x*_*kj*_ ‖^2^ in Equation 17 by *RF*(*T*_*ki*_, *T*_*kj*_) in order to obtain the approximation formula for *SS*_*W*_:

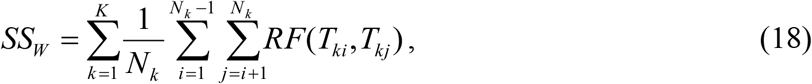

where *T*_*ki*_ and *T*_*kj*_ are trees *i* and *j* of cluster *k*, respectively.

Also, in the case of the Euclidean distance we have:

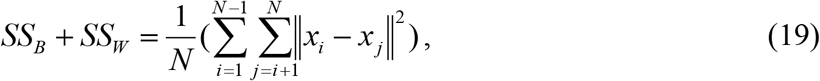

where *x*_*i*_ and *x* _*j*_ are two different objects in *X* (Caliński and Harabasz 1974).

Thus, the approximation of the global variance between groups, *SS*_*B*_, can be calculated as follows:

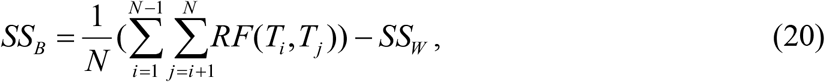

where *T*_*i*_ and *T*_*j*_ are trees *i* and *j* in a given set of trees Π, and *SS*_*W*_ is calculated according to Equation 18.

As the square root of the Robinson and Foulds distance has the Euclidean properties, Equations 18 and 20 establish the exact formulas for calculating the indices *SS*_*B*_ and *SS*_*W*_ for the objective function *OF*_*EA*_ defined by Equation 2. Obviously, the objective function *OF*_*EA*_ is only an approximation of the objective function defined in Equation 1 because the centroid of a cluster of trees is not necessarily a consensus tree of the cluster. Moreover, it is not necessarily a tree. However, as we show it in the Results section (see the main text), this approximation provides very good classification results when clustering trees.

#### Silhouette index adapted for tree clustering

Another popular index considered in our study is the Silhouette width (*SH*) (Rousseeuw 1987). Traditionally, the Silhouette width of cluster *k* is defined as follows:

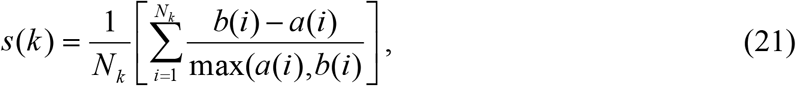

where *N*_*k*_ is the number of objects belonging to cluster *k, a*(*i*) is the average distance between object *i* and all other objects belonging to cluster *k*, and *b*(*i*) is the smallest, over all clusters *k*’ different from *k*, of all average distances between *i* and all the objects of cluster *k’*.

We used Equations 22 and 23 for calculating *a*(*i*) and *b*(*i*), respectively, in our tree clustering algorithm (see also Tahiri et al. 2018). For instance, the quantity *a*(*i*) can be determined as follows:

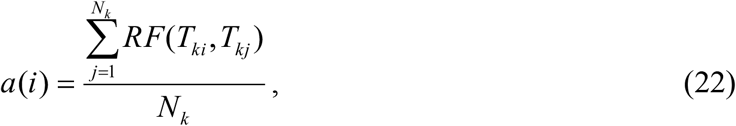

and the formula for *b*(*i*) is as follows:

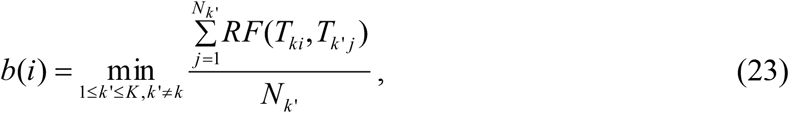

where *T*_*k’j*_ is tree *j* of cluster *k’*, such that *k’* ≠ *k*, and *N*_*k’*_ is the number of trees in cluster *k’*. The optimal number of clusters, *K*, corresponds to the maximum average Silhouette width, *SH*, which is calculated as follows:

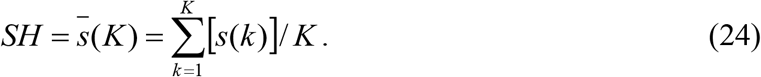

The value of the Silhouette index defined by Equation 24 is located between -1 and +1.

#### Gap statistic adapted for tree clustering

Unfortunately, the *CH* and *SH* cluster validity indices defined by Equations 15 and 24 do not allow us to compare the solution consisting of a single consensus tree (*K* = 1; the calculation of *CH* and *SH* is impossible in this case) with clustering solutions involving multiple consensus trees (*K* ≥ 2). This can be viewed as an important drawback of the *CH* and *SH*-based classifications because a good tree clustering method should be able to recover a single consensus tree when the input set of trees is homogeneous (e.g. in case of gene trees that share the same evolutionary history).

The *Gap* statistic was first used by Tibshirani et al. (2001) to estimate the number of clusters provided by partitioning algorithms. The formulas proposed by Tibshirani et al. were based on the properties of the Euclidean distance. In the context of tree clustering, the *Gap* statistic can be defined as follows. Consider a clustering of *N* trees into *K* non-empty clusters, where *K* ≥ 1. We first define the total intracluster distance, *D*_*k*_, characterizing the cohesion between the trees belonging to the same cluster *k*:

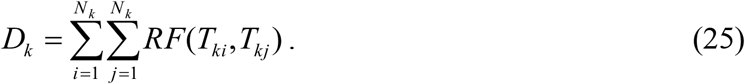

Then, the sum of the average total intracluster distances, *V*_*K*_, can be calculated:

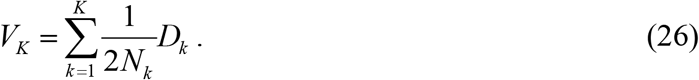

Finally, the *Gap* statistic, which reflects the quality of a given clustering solution with *K* clusters, can be defined as follows:

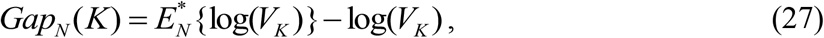

where ^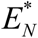^ denotes expectation under a sample of size *N* from the reference distribution. The following formula (Tibshirani et al. 2001) for the expectation of log(*V*_*K*_) was used in our method:

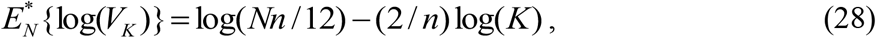

where *n* is the number of tree leaves.

The largest value of the *Gap* statistic corresponds to the best clustering.

#### Ball-Hall index adapted for tree clustering

Ball and Hall (1965) introduced the ISODATA procedure to measure the average dispersion of groups of objects with respect to the mean square root distance, i.e. the intra-group distance, which would lead to the following formula in case of tree clustering:

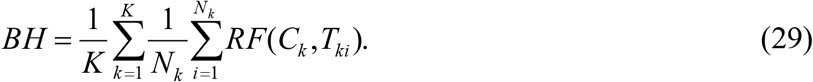

Replacing the inner sum of Equation 29 by its Euclidean approximation (as in Equation 2), we obtain the following formula:

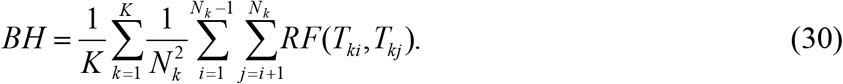

### A.3 Theorem establishing the lower and the upper bounds of the objective function *OF*

#### Theorem 1

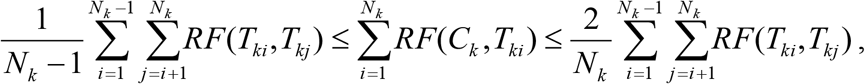

*where N*_*k*_ *is the number of trees in cluster k, T* _*ki*_ *and T*_*kj*_ *are, respectively, trees i and j in cluster k, and C*_*k*_ *is the majority-rule consensus tree of cluster k*.

*Proof*

First, the sum 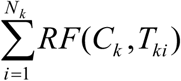 can be decomposed into the following double sum:

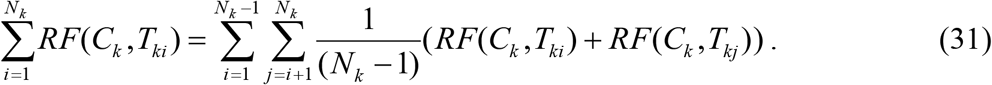

We know that the Robinson and Foulds distance is a metric, and thus satisfies the triangular inequality (Robinson and Foulds 1981). Hence, the following inequality holds for any pair of trees (*T*_*ki*_,*T*_*kj*_): *RF*(*C*_*k*_, *T*_*ki*_) + *RF*(*C*_*k*_, *T*_*kj*_) ≥ *RF*(*T*_*ki*_, *T*_*kj*_).

This means that:

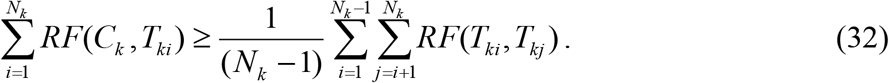

Second, based on the property, proved by Barthélemy and McMorris, that the majority consensus tree of a set of trees is a median tree of this set in the sense of the *RF* distance (Barthélemy and McMorris 1986; Barthélemy and Monjardet 1981), we have: 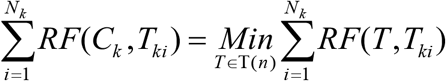, where T(*n*) is the set of all possible phylogenetic trees with *n* leaves.

Thus, we obtain:

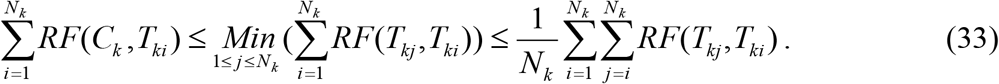

It is easy to see that the upper bound in Equation 33 equals to: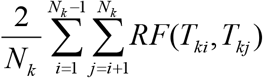 Obviously, the term 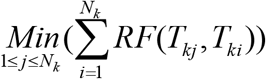 can be also used as an upper bound of 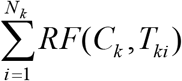. □

### A.4 Detailed simulation protocol

This section present the detailed simulation protocol adopted in our study. The data generation procedure used in the first experiment (i.e. with multiple consensus trees) included three main steps. First, we randomly generated a species phylogeny *T*_0_ with *n* leaves (i.e. it played the role of the first consensus tree in our simulations) using the HybridSim program of Woodhams et al. (2016). Second, still using HybridSim, we generated *K*-1 gene phylogenies, *T*_1_, …, *T*_*K*_, defined on the same set of *n* leaves (i.e. they played the role of the other consensus trees in our simulations). Each of these phylogenies differed from *T*_0_ by a specific number of hybridization events (the value of the *hybridization_rate* parameter in HybridSim varied from 1 to 4 in our experiments; this value was drawn randomly using a uniform distribution). The number of clusters, *K*, in this experiment varied from 1 to 5, and the number of tree leaves, *n*, was taking the values 8, 16, 32, 64 (and 128, when the comparison with the state-of-the-art tree partitioning algorithms was carried out). The HybridSim program allows one to generate phylogenies in the presence of hybridization and coalescence/incomplete lineage sorting events. Thus, we used the *hybridization_rate* parameter of HybridSim to generate centers of gene tree clusters (i.e. multiple consensus trees or multiple supertrees) and the *coalescence_rate* parameter to generate incongruence across gene trees. The value of the *coalescence_rate* parameter, adding some noise to gene phylogenies, varied between 1 (high noise) and 10 (low noise); the other HybridSim parameters were the default program parameters. Third, for each gene phylogeny *T*_*i*_ (*i* = 1, …, *K*), being the center of cluster *i*, we randomly generated a set of 100 trees *T*_*i*_’ (the number of trees *T*_*i*_’ varied form 20 to 100 with the step of 20 in the simulation conducted with the state -of-the-art tree partitioning methods; see Fig. 2 in the main text), belonging to cluster *i*, such that each tree *T*_*i*_’ differed from *T*_*i*_ by a specific number of coalescence/incomplete lineage sorting patterns of incongruence, which was controlled through the value of the *coalescence_rate* parameter.

First, we compared the classification performances (in terms of Adjusted Rand Index, ARI), of the six variants of our tree clustering algorithm applied to trees with identical sets of leaves (see Fig. 1). The six evaluated variants of our algorithm were based on: (1) the *OF*_*EA*_ objective function with approximation by Euclidean distance and *CH* cluster validity index (*CH-E*), (2) the *OF*_*LA*_ objective function with approximation by the lower bound and *CH* cluster validity index (*CH-LB*), (3) the *OF*_*MA*_ objective function with approximation by the mean of interval and *CH* cluster validity index (*CH-MI*), (4) the *OF*_*EA*_ objective function and Silhouette cluster validity index (*SH*), (5) the *OF*_*EA*_ objective function and the Gap cluster validity index (*Gap*), and (6) *OF*_*EA*_ objective function and Ball and Hall cluster validity index (*BH*).

Second, the variant of our algorithm based on the *OF*_*EA*_ objective function with approximation by Euclidean distance and *CH* cluster validity index (*CH-E*) that showed the best overall performance in the first simulation was compared to the state-of-the-art tree clustering methods, including: (1) the MPC method by Bonnard et al. (2006), (2) the tree clustering algorithm by Stockham et al. (2002), which is based on *k*-means clustering with squared *RF* distance (this method recomputes the majority-rule consensus trees of all clusters at each *k*-means iteration), and (3) the *k*-medoids tree clustering algorithm by Tahiri et al. (2018), which uses the *RF* distance and *SH* cluster validity index. The comparison was conducted in terms of the quality of clustering results returned by competing methods (Fig. 2a and b) and the running time (Fig. 2c and d).

Our second simulation experiment involved partitioning trees with different sets of leaves with the objective to build multiple consensus supertrees. The data generation protocol for this experiment included an additional step consisting of the random removal of some species (i.e. tree leaves) from the generated gene trees. The branches adjacent to the removed leaves were collapsed. The following intervals of missing data were considered: 0% (no species were removed), 10% (5% to 15% of species were randomly removed), 25% (16% to 35% of species were randomly removed) and 50% (36% to 65% of species were randomly removed). The exact number of species to be removed from each gene tree and each data interval was drawn randomly using a uniform distribution. We also made sure that every pair of trees in each input dataset had at least 4 species in common. The value of the penalization parameter *α* in Equation 10 was set to 0 in this experiment. Two independent simulations were carried for the supertree version of our algorithm using the *OF*_*ST_EA*_ objective function (Equation10) and the *CH-E* (Equations 11-12) and *BH* cluster validity indices adapted for supertree partitioning (see Figs A3 and A4). The *OF*_*ST_EA*_ objective function was used because it provided the best overall performance in our first simulation experiment with consensus trees (see Fig. 1), whereas the *BH* index was used because it slightly outperformed the *Gap* index in case of heterogeneous data (i.e. when the number of clusters *K* was equal to 1).

Our third simulation experiment was also conducted to evaluate the ability of our algorithm to cope with incomplete data. As in the second experiment, gene trees with different sets of leaves were considered. The supertree version of our algorithm based on the *OF*_*ST_EA*_ objective function (Equation 10) and the *CH-E* cluster validity index (Equations 11-12) was used here with different values of the penalization parameter *α*, which varied from 0 to 1 (with the step of 0.2; see Fig. 3).

In all simulation experiments, our tree partitioning algorithm was carried out with 100 random starts until the convergence of the selected objective function or until 50 iterations in the algorithm’s internal loop were completed (i.e. the same stopping rule as in the traditional *k*-means algorithm were applied). All reported ARI results (see Figs 1 to 3 and A3-A4) are the averages taken over all considered numbers of trees, leaves and clusters. The simulation results presented in Figures 1, A3 and A4 (portions a, b and d, in all these figures) and Figures 2 and 3 correspond to the case where the value of the coalescence rate parameter was fixed to 5. Figures 1, A3 and A4 (portion c, in all these figures) illustrate how the algorithm’s results vary with respect to the change in the coalescence rate.

Our computational experiments were carried out using a 64-bit PC computer equipped with an Intel i7-8750H CPU (2.5 GHz) and 32 Gb of RAM, except for the simulation comparing the performances of the state-of-the-art clustering algorithms, which was conducted on a high-performance parallel computing server of Compute Canada.

### A.5 Simulations with multiple alternative supertrees using *CH* and *BH* indices

Figures A3 and A4 illustrate the classification performance of our supertree clustering algorithm based on the *OF*_*ST_EA*_ objective function used with *CH* and *BH* cluster validity indices, respectively. Here, the value of the penalization parameter *α* in Equation 10 was set to 0.

The algorithm was applied to gene trees containing different numbers and sets of leaves. We can observe that the clustering performances of both tested algorithm’s variants gradually decreases as the number of missing species (i.e. tree leaves) increases. The supertree clustering algorithm based on *CH* generally outperformed that based on *BH*, but the *BH*-based variant seems to be less sensitive to the increase in the number of missing species as the number of clusters and the coalescence noise grow (e.g. for the case of 5 tree clusters, the average ARI value provided by the *CH*-based version decreased from 0.97 for 0% of missing species to 0.81 for 50% of missing species (Fig. A3a), whereas for the *BH*-based version it decreased from 0.51 for 0% of missing species to 0.44 for 50% of missing species (Fig. A4a)).

**Fig. A3.**
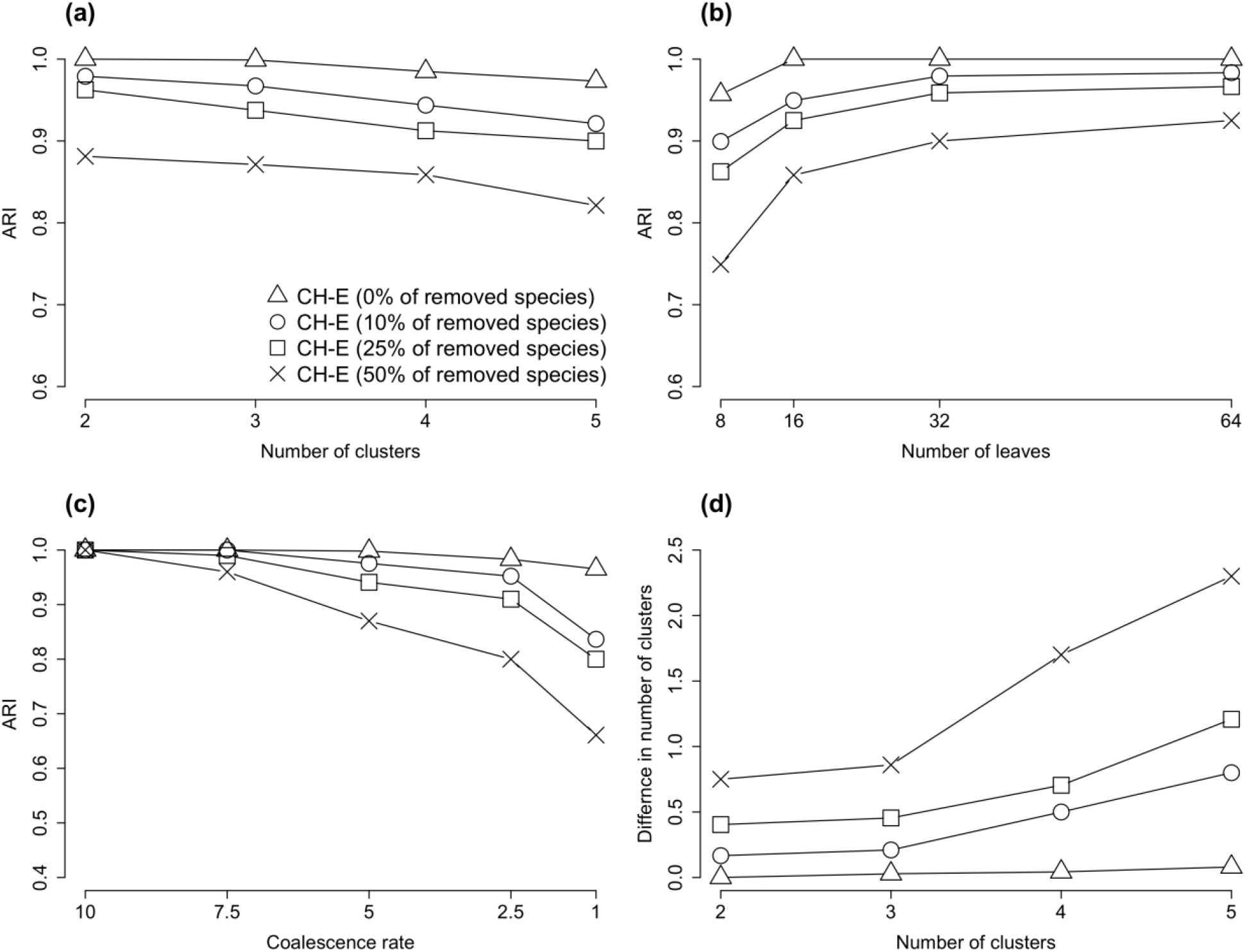
Classification performance of our *k*-means-based supertree clustering algorithm applied to trees with different numbers and sets of leaves (the following numbers of leaves were randomly removed from each tree: 0% (no species were removed), 10% (5% to 15% of species were removed), 25% (16% to 35% of species were removed) and 50% (36% to 65% of species were removed)) using the *OF*_*ST_EA*_ objective function with approximation by Euclidean distance and the *CH* cluster validity index (*CH-E*). The value of the penalization parameter *α* was set to 0. The results are presented in terms of *ARI* with respect to the: (a) number of tree clusters, (b) number of leaves and (c) coalescence rate; (d) average absolute difference between the true and the obtained number of clusters. The coalescence rate parameter in the HybridSim program was set to 5 in simulations (a), (b) and (d). The presented results are the averages taken over all considered numbers of leaves in simulations (a), (c) and (d), and all considered numbers of clusters in simulations (b) and (c).

**Fig. A4.**
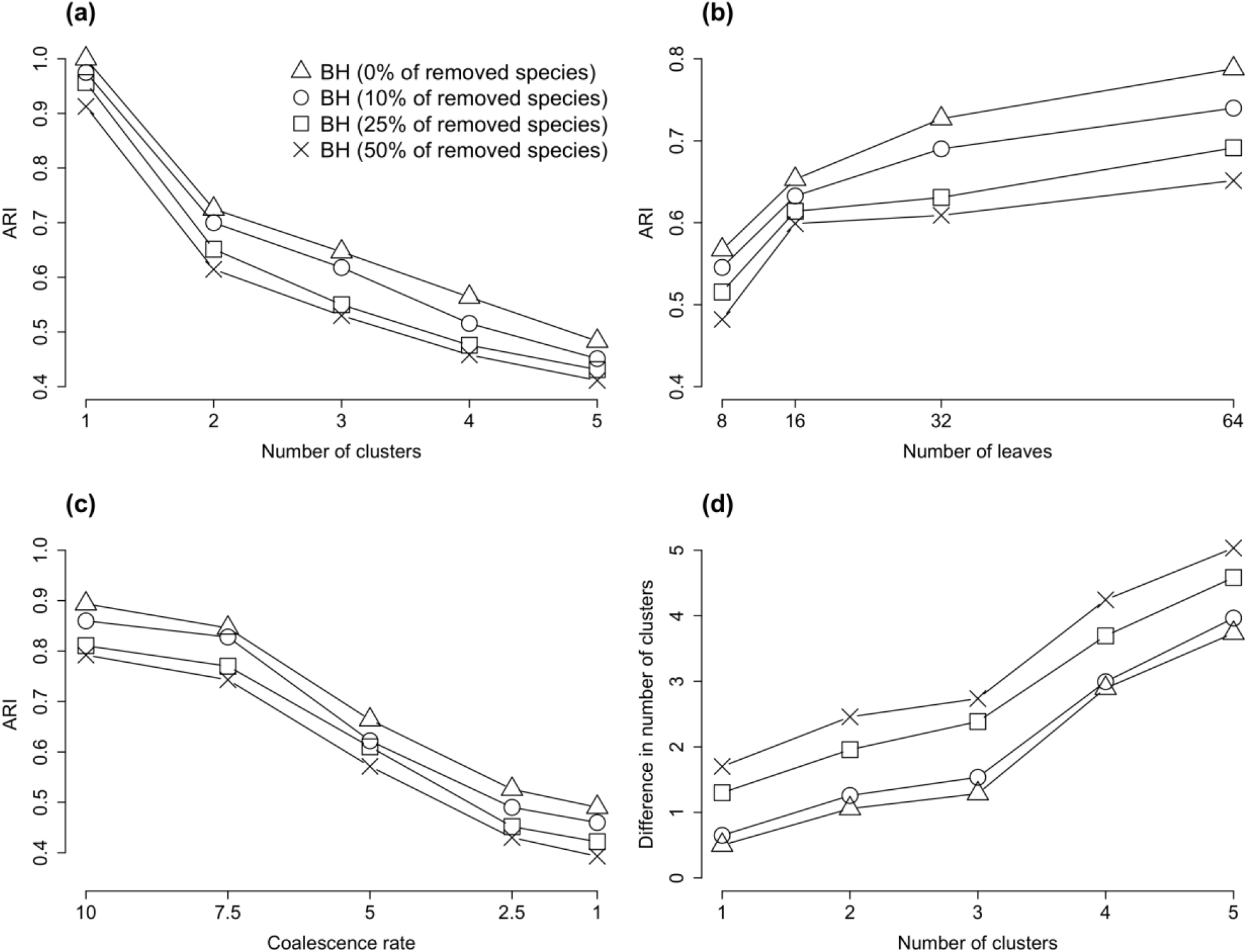
Classification performance of our *k*-means-based supertree clustering algorithm applied to trees with different numbers and sets of leaves using the *OF*_*ST_EA*_ objective function with approximation by Euclidean distance and the Ball and Hall cluster validity index (*BH*). Figure A3 panel description applies here.

## Supplementary Material

**Supplementary Table 1.** The full virus names, abbreviations, host species and GenBank/GISAID accession numbers for the 43 betacoronavirus genomes analysed in our study.

| Organism's name | Abbreviation | Host | Accession number (GenBank, GISAID) |
| --- | --- | --- | --- |
| hCoV-19/Australia/VIC231/2020 | Hu Australia VIC231 2020 | Human | EPI_ISL_419 926 |
| hCoV-19/USA/UT-00346/2020 | Hu USA UT 00346 20202 | Human | EPI_ISL_420 819 |
| BetaCoV/Wuhan-Hu-1 | Hu Wuhan 2020 | Human | NC_045512.2 |
| hCoV-19/Italy/TE4836/2020 | Hu Italy TE4836 2020 | Human | EPI_ISL_418 260 |
| Bat coronavirus RaTG13 | RaTG13 | Bat | MN996532.1 |
| hCoV-19/pangolin/Guangdong/1/2019 | Guangdong Pangolin 1 2019 | Pangolin | EPI_ISL_410 721 |
| hCoV-19/pangolin/Guangdong/P2S/2019 | Guangdong Pangolin P2S 2019 | Pangolin | EPI_ISL_410 544 |
| PCoV_GX-P5E | Guangxi Pangolin P5E | Pangolin | MT040336 |
| PCoV_GX-P2V | Guangxi Pangolin P2V | Pangolin | MT072864 |
| PCoV_GX-P5L | Guangxi Pangolin P5L | Pangolin | MT040335 |
| PCoV_GX-P1E | Guangxi Pangolin P1E | Pangolin | MT040334.1 |
| PCoV_GX-P4L | Guangxi Pangolin P4L | Pangolin | MT040333 |
| bat-SL-CoVZC45 | Bat CoVZC45 | Bat | MG772933.1 |
| bat-SL-CoVZXC21 | Bat CoVZXC21 | Bat | MG772934.1 |
| BtCoV/273/2005 | BtCoV 273 2005 | Bat | DQ648856.1 |
| Bat SARS coronavirus Rf1 | Rf1 | Bat | DQ412042.1 |
| Bat SARS coronavirus HKU3-12 | HKU3-12 | Bat | GQ153547.1 |
| Bat SARS coronavirus HKU3-6 | HKU3-6 | Bat | GQ153541.1 |
| BtCoV/279/2005 | BtCoV 279 2005 | Bat | DQ648857.1 |
| SARS coronavirus BJ01 | SARS | Human | AY278488.2 |
| SARS coronavirus | Tor2 | Human | NC_004718.3 |
| SARS coronavirus BJ182-4 | SARS-CoV BJ182-4 | Human | EU371562 |
| Bat SARS-like coronavirus<br>Rs3367 | Rs3367 | Bat | KC881006.1 |
| SARS-related coronavirus<br>BtKY72 | BtKY72 | Bat | KY352407.1 |
| Bat coronavirus BM48-31/BGR/2008 | BM48 31 BGR 2008 | Bat | GU190215.1 |
| Human betacoronavirus 2c<br>EMC/2012 | MERS-CoV S | Human | JX869059 |
| Bat coronavirus HKU5-1 | Bat HKU5-1 | Bat | NC_009020 |
| Bat coronavirus HKU4-1 | Bat HKU4-1 | Bat | NC_009019 |
| Feline infectious peritonitis virus | Feline per | Cat | NC_002306 |
| Human coronavirus HKU1 | HKU1 | Human | NC_006577 |
| Murine coronavirus RA59/R13 | Murine RA59/R13 | Mouse | ACN89689 |
| Murine hepatitis virus strain 4 | Murine hep 4 | Mouse | P22432 |
| Mouse hepatitis virus strain<br>MHV-A59 C12 mutant | Murine A59 | Mouse | NC_001846 |
| Murine hepatitis virus | Murine virus | Mouse | ABS87264 |
| Rat coronavirus Parker | Rat Parker | Rat | NC_012936 |
| Rabbit coronavirus HKU14 | Rabbit HKU14 | Rabbit | NC_017083 |
| Equine coronavirus | Equine NC99 | Horse | NC_010327 |
| Porcine hemagglutinating en-<br>cephalomyelitis virus | Porcine v | Pig | NC_007732 |
| Human coronavirus OC43 | Human OC43 | Human | NC_005147 |
| Human enteric coronavirus strain<br>4408 | Human ent 4408 | Human | NC_012950 |
| Bovine coronavirus | Bovine CoV | Calf | NC_003045 |
| Bovine respiratory coronavirus<br>AH187 | Bovine AH187 | Calf | NC_012948 |
| Bovine respiratory coronavirus<br>bovine/US/OH-440-TC/1996 | Bovine OH440 | Calf | NC_012949 |

### Tree for gene ORF1ab (43 taxa)

**Supplementary Figure 1.**
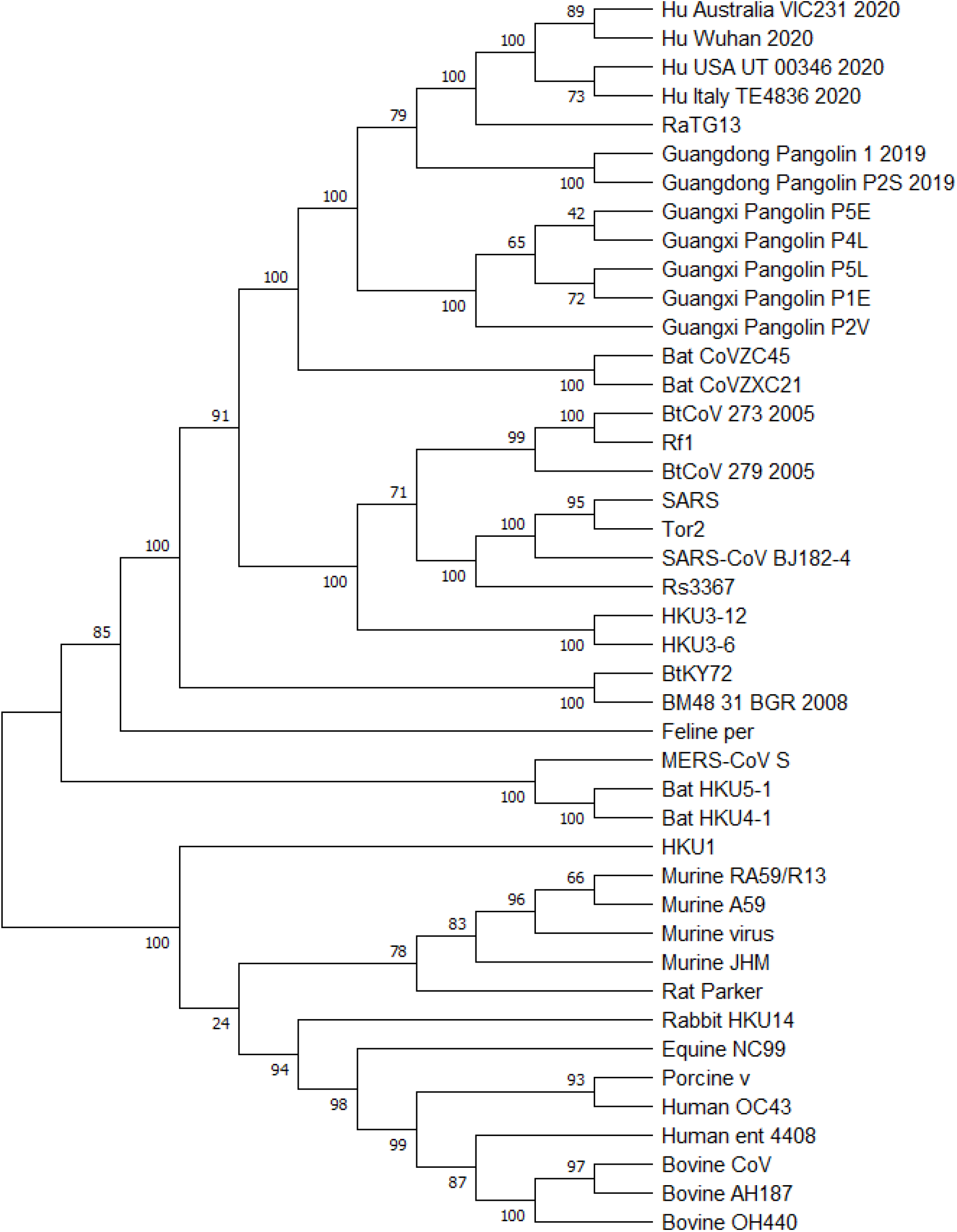
Phylogenetic tree of gene ORF1ab inferred using RAxML with 100 bootstrap replicates for a group of 43 betacoronaviruses (see Supplementary Table 1).

### Tree for gene S (43 taxa)

**Supplementary Figure 2.**
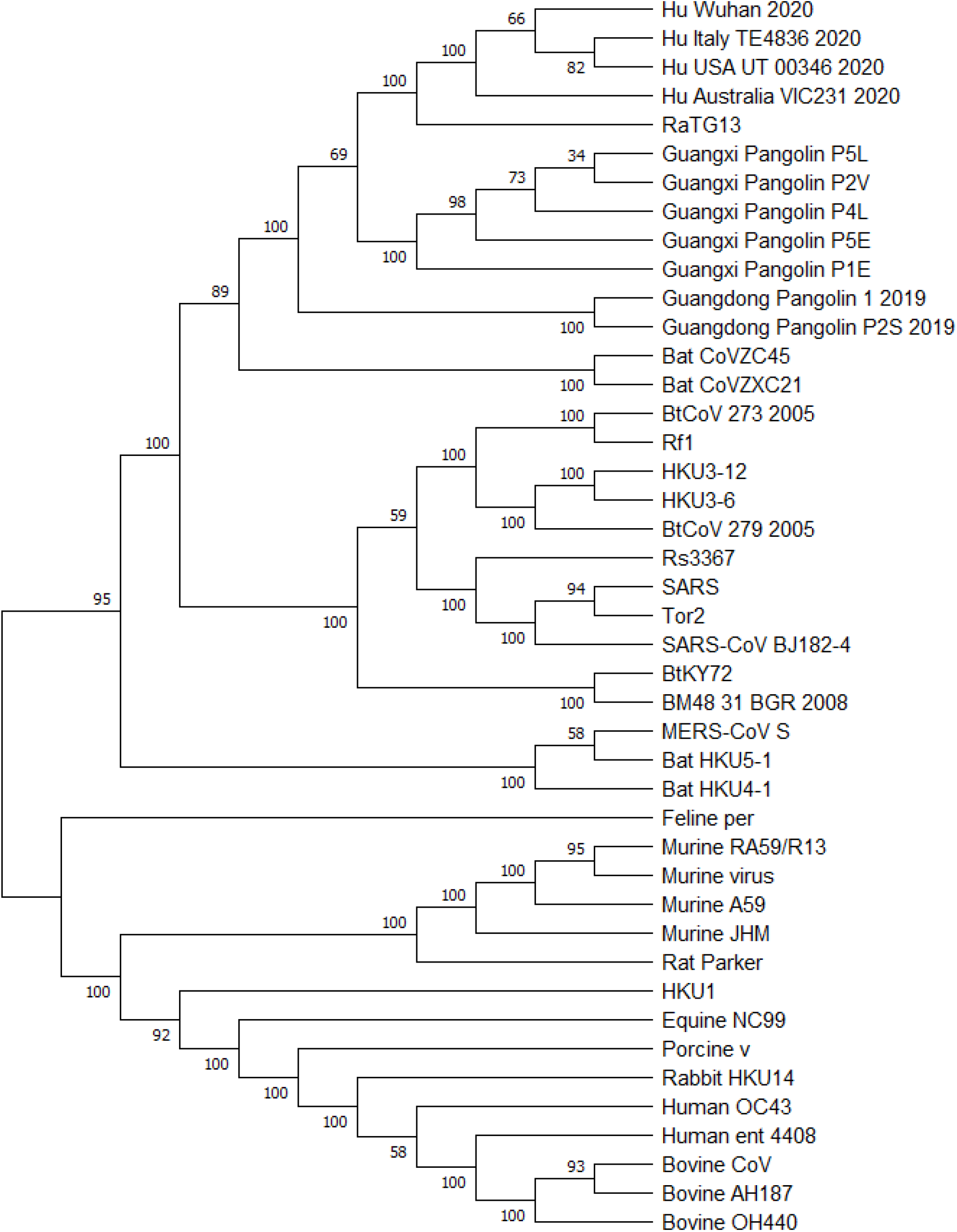
Phylogenetic tree of gene S inferred using RAxML with 100 boo t-strap replicates for a group of 43 betacoronaviruses (see Supplementary Table 1).

### Tree for gene ORF3a (43 taxa)

**Supplementary Figure 3.**
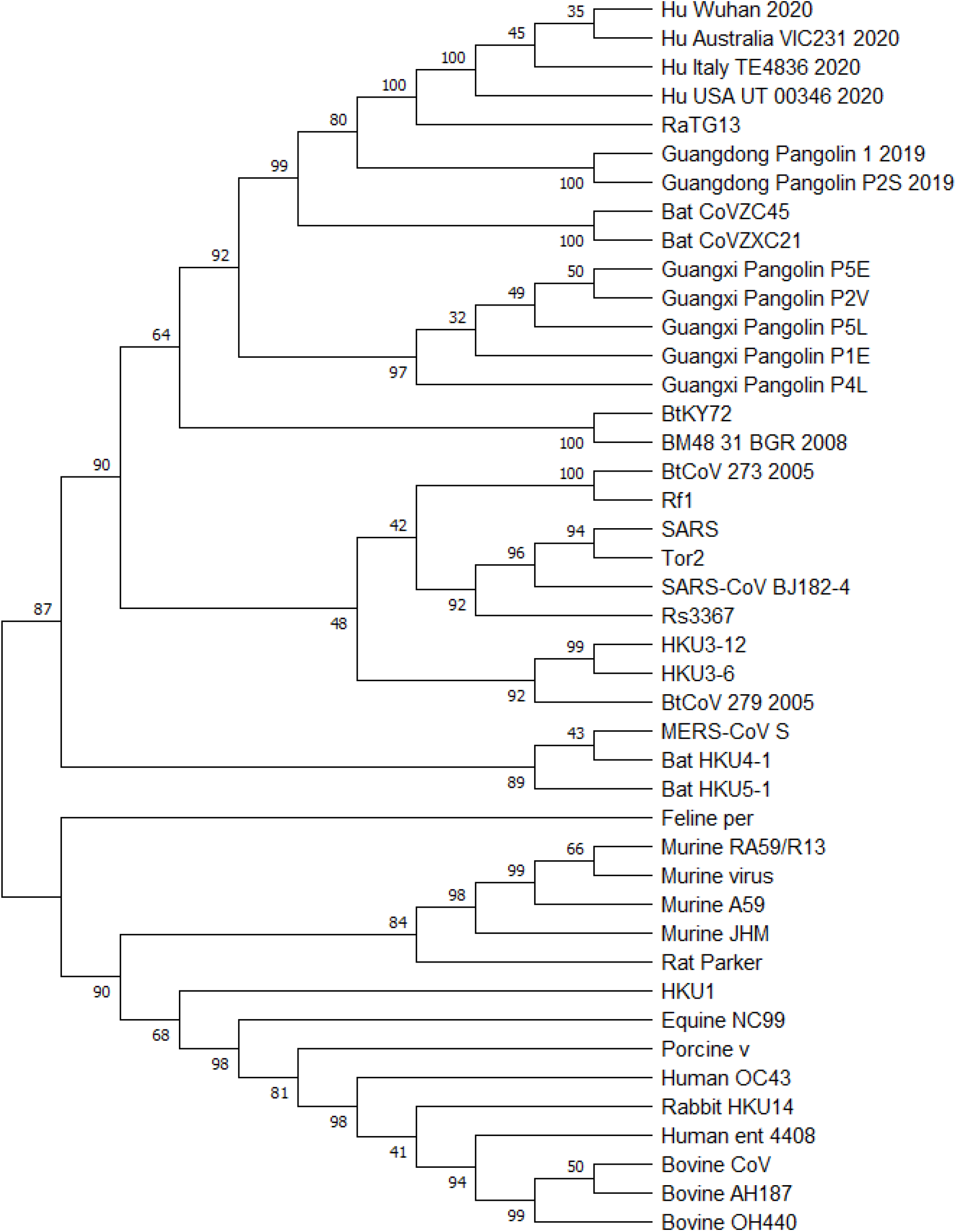
Phylogenetic tree of gene ORF3a inferred using RAxML with 100 bootstrap replicates for a group of 43 betacoronaviruses (see Supplementary Table 1).

### Tree for gene E (43 taxa)

**Supplementary Figure 4.**
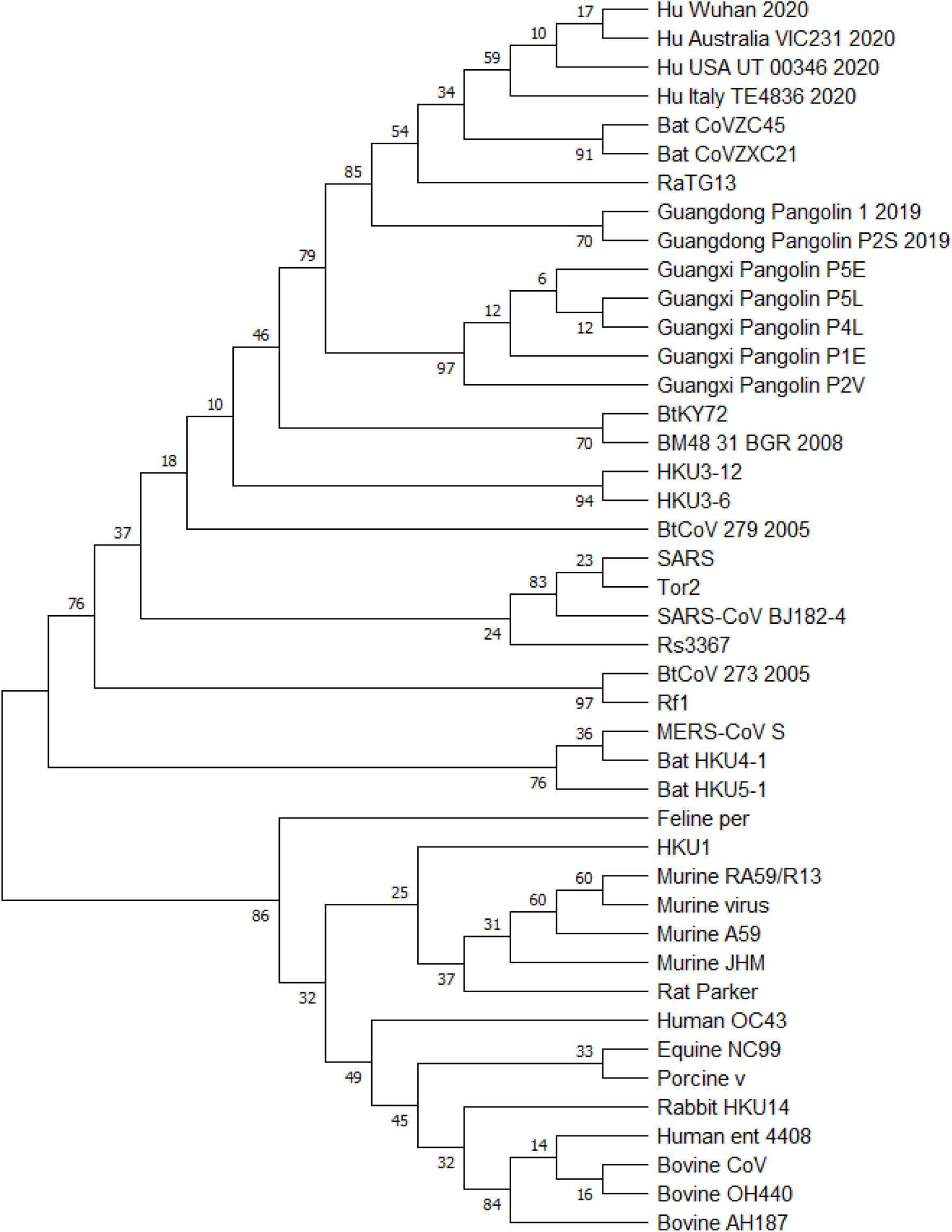
Phylogenetic tree of gene E inferred using RAxML with 100 boo t-strap replicates for a group of 43 betacoronaviruses (see Supplementary Table 1).

### Tree for gene M (43 taxa)

**Supplementary Figure 5.**
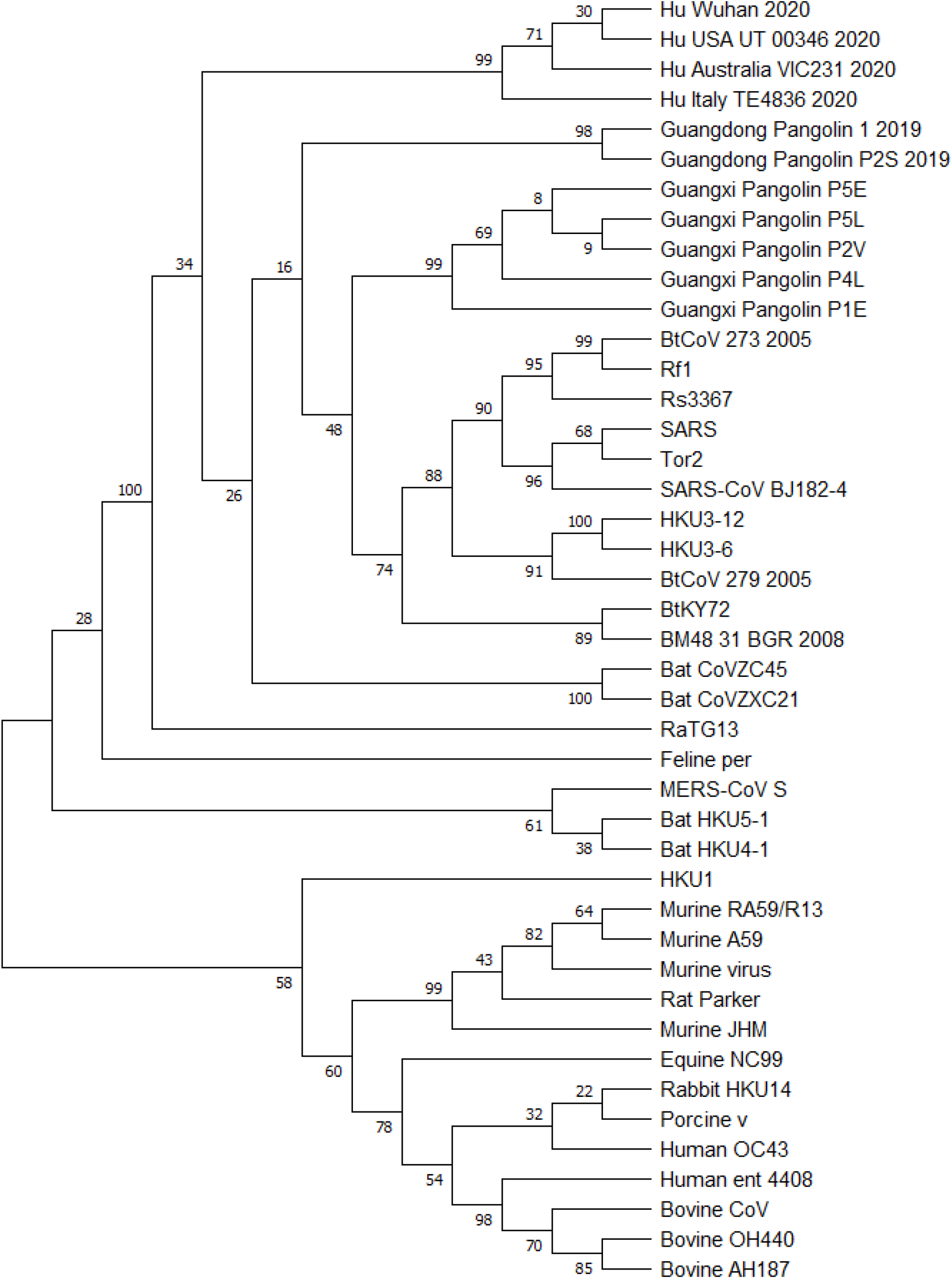
Phylogenetic tree of gene M inferred using RAxML with 100 bootstrap replicates for a group of 43 betacoronaviruses (see Supplementary Table 1).

### Tree for gene ORF6 (25 taxa)

**Supplementary Figure 6.**
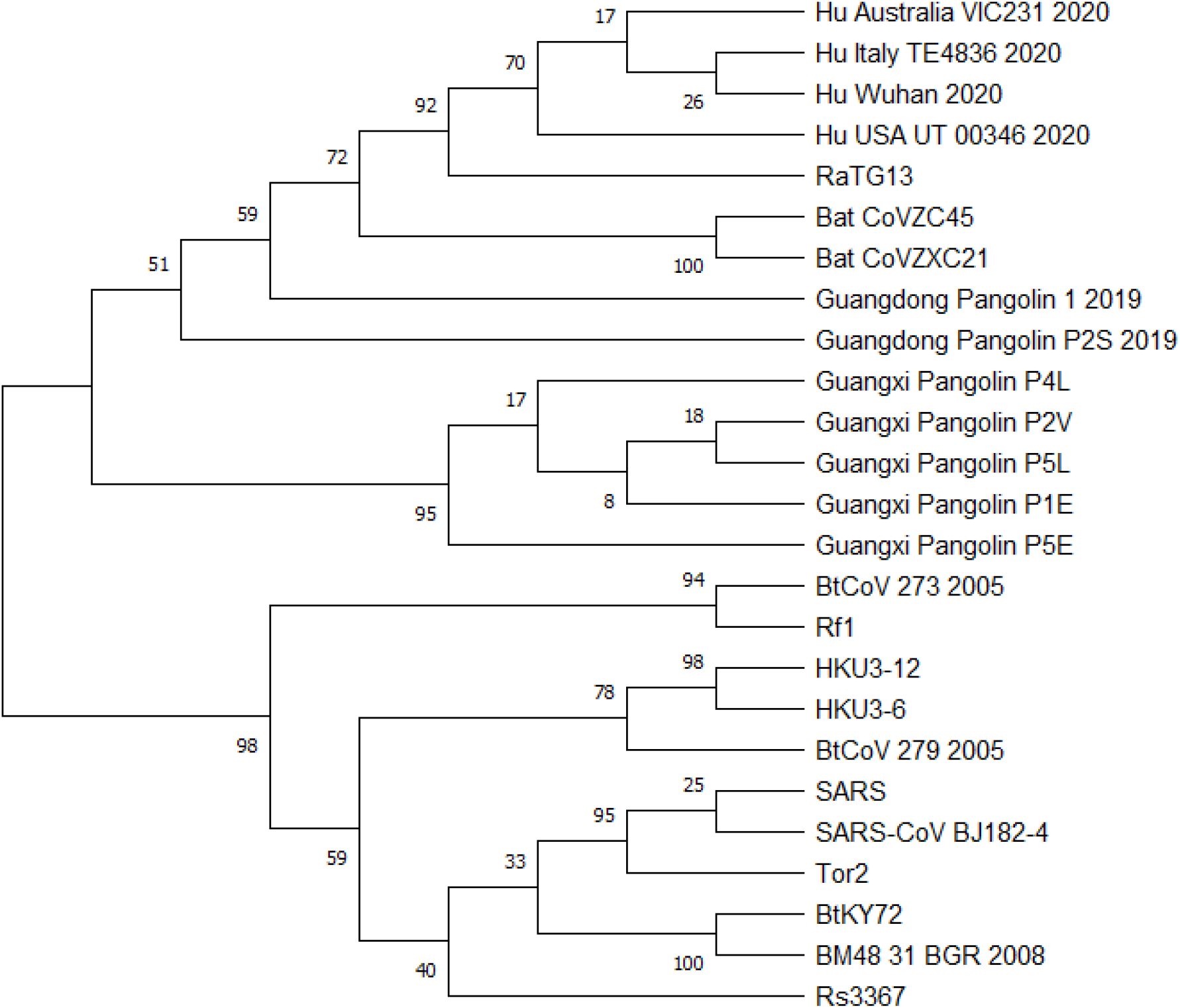
Phylogenetic tree of gene ORF6 inferred using RAxML with 100 bootstrap replicates for a group of 25 betacoronaviruses (see Supplementary Table 1).

### Tree for gene ORF7a (25 taxa)

**Supplementary Figure 7.**
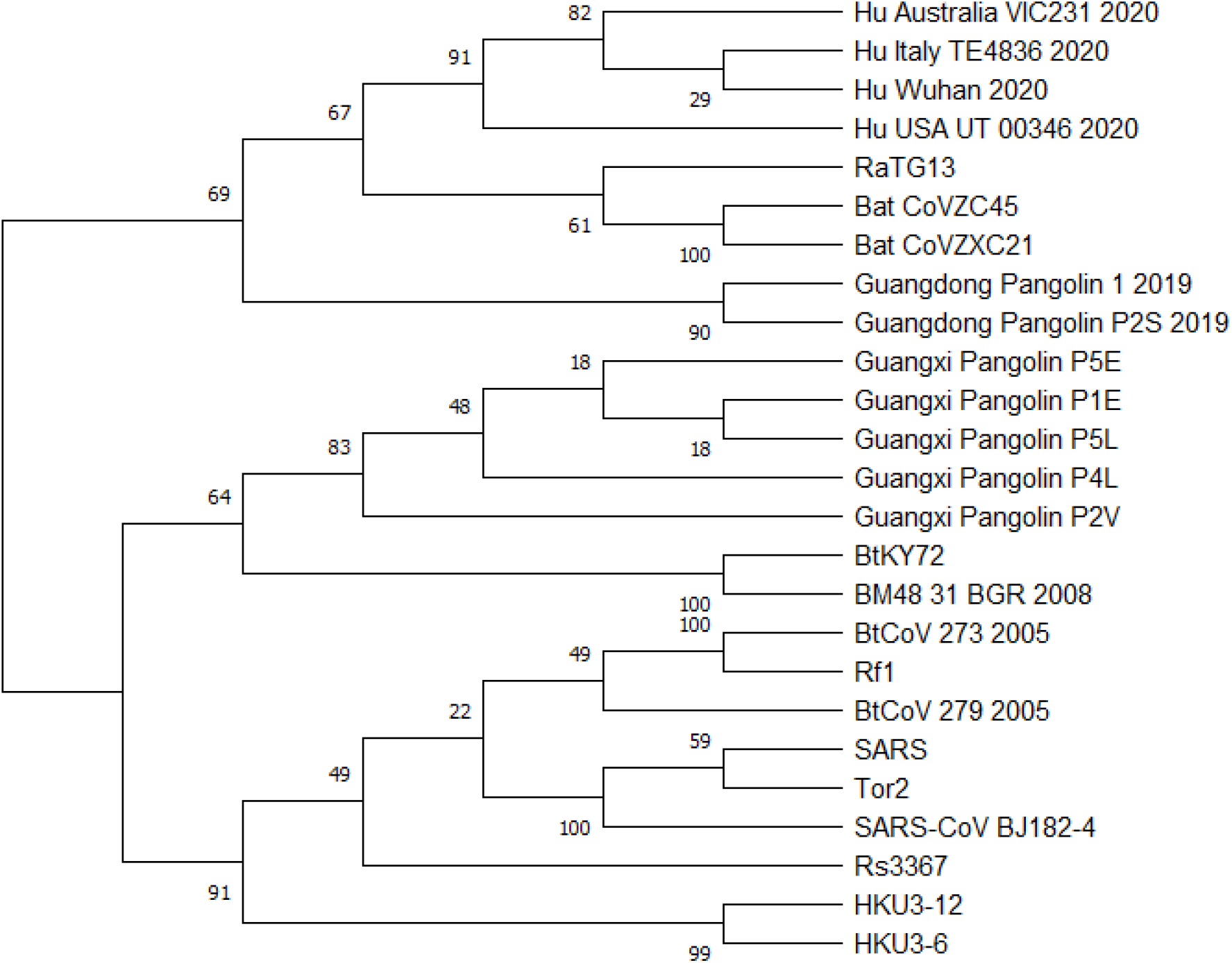
Phylogenetic tree of gene ORF7a inferred using RAxML with 100 bootstrap replicates for a group of 25 betacoronaviruses (see Supplementary Table 1).

### Tree for gene ORF7b (25 taxa)

**Supplementary Figure 8.**
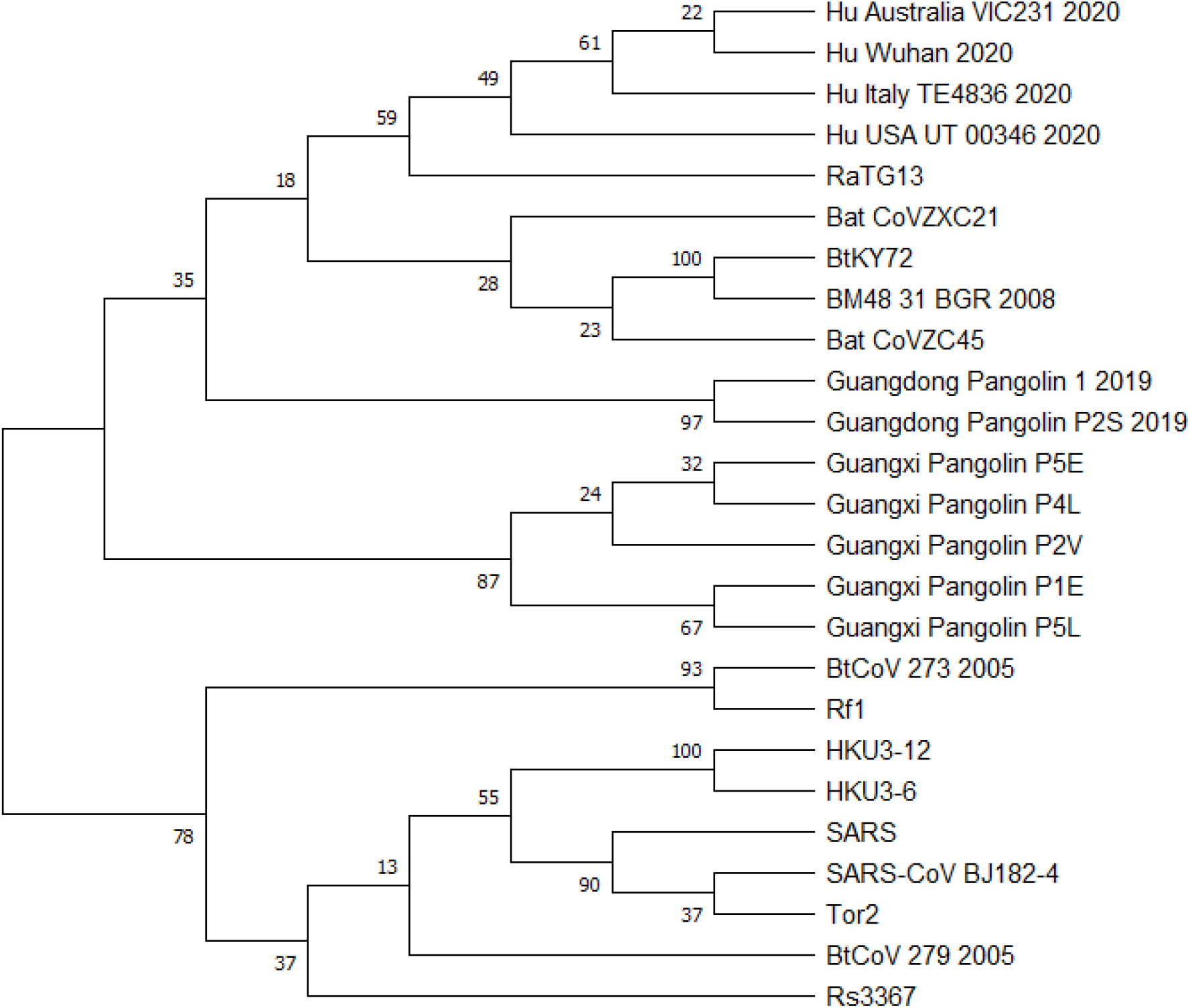
Phylogenetic tree of gene ORF7b inferred using RAxML with 100 bootstrap replicates for a group of 25 betacoronaviruses (see Supplementary Table 1).

### Tree for gene ORF8 (23 taxa)

**Supplementary Figure 9.**
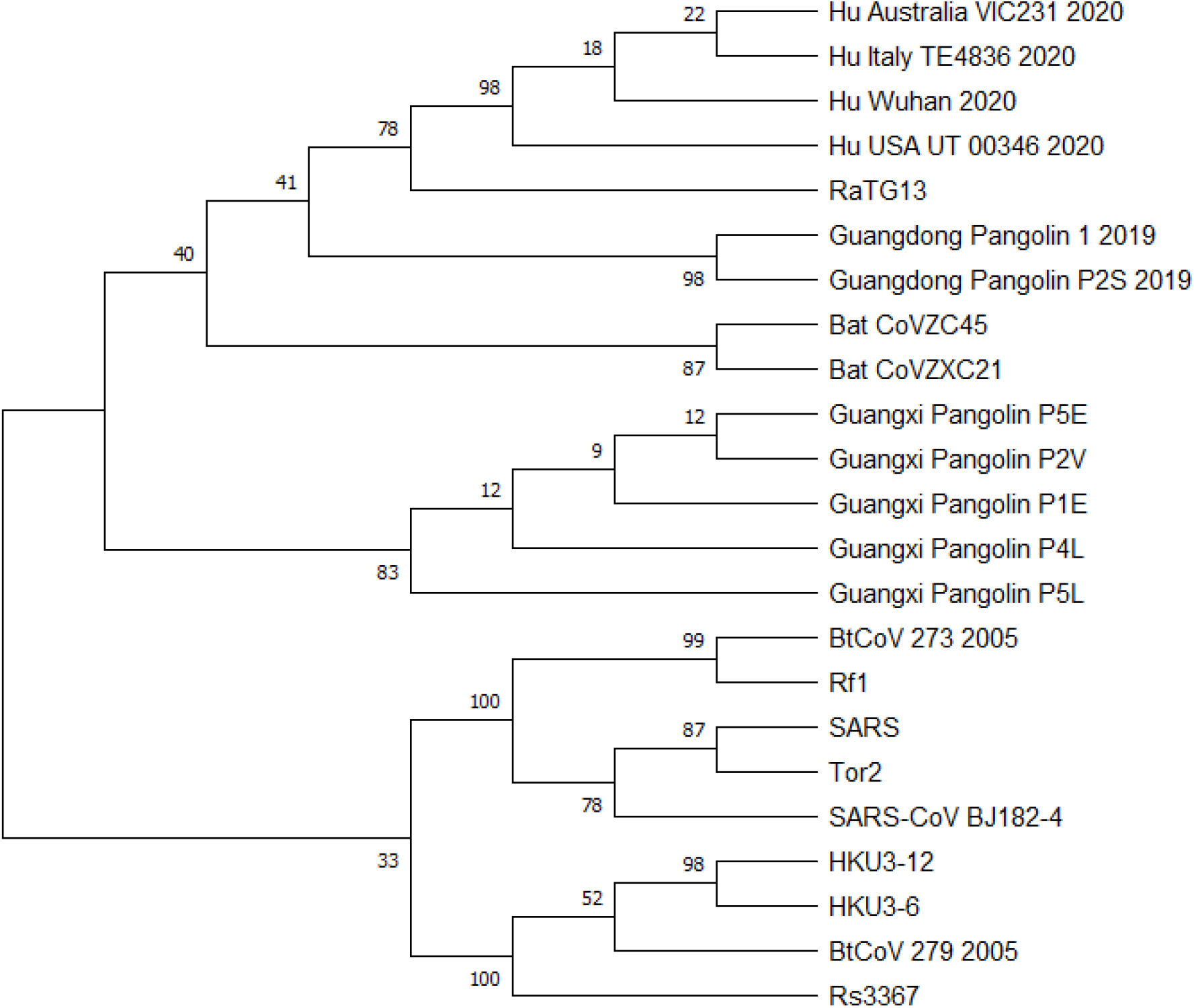
Phylogenetic tree of gene ORF8 inferred using RAxML with 100 bootstrap replicates for a group of 23 betacoronaviruses (see Supplementary Table 1).

### Tree for gene N (43 taxa)

**Supplementary Figure 10.**
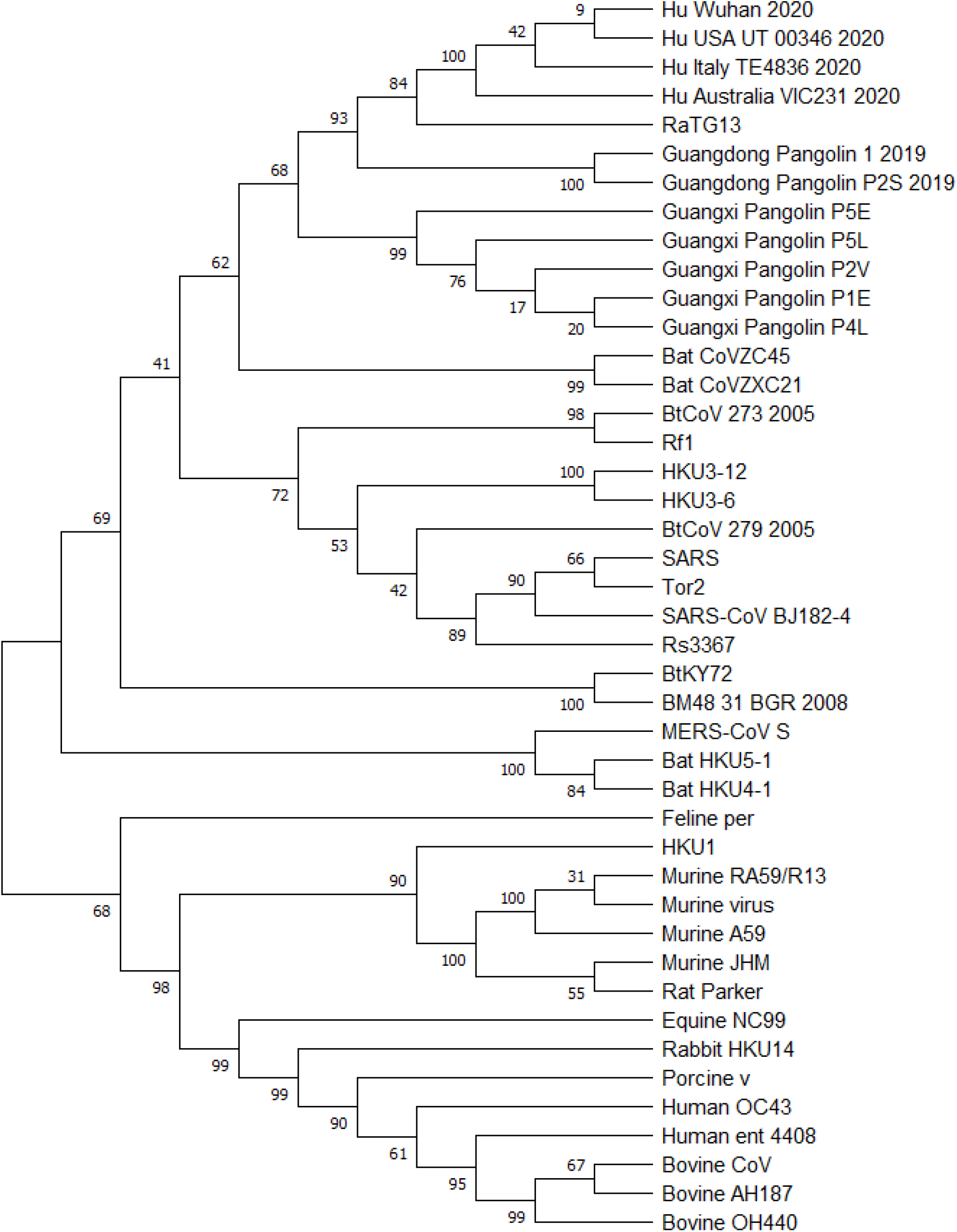
Phylogenetic tree of gene N inferred using RAxML with 100 bootstrap replicates for a group of 43 betacoronaviruses (see Supplementary Table 1).

### Tree for gene ORF10 (25 taxa)

**Supplementary Figure 11.**
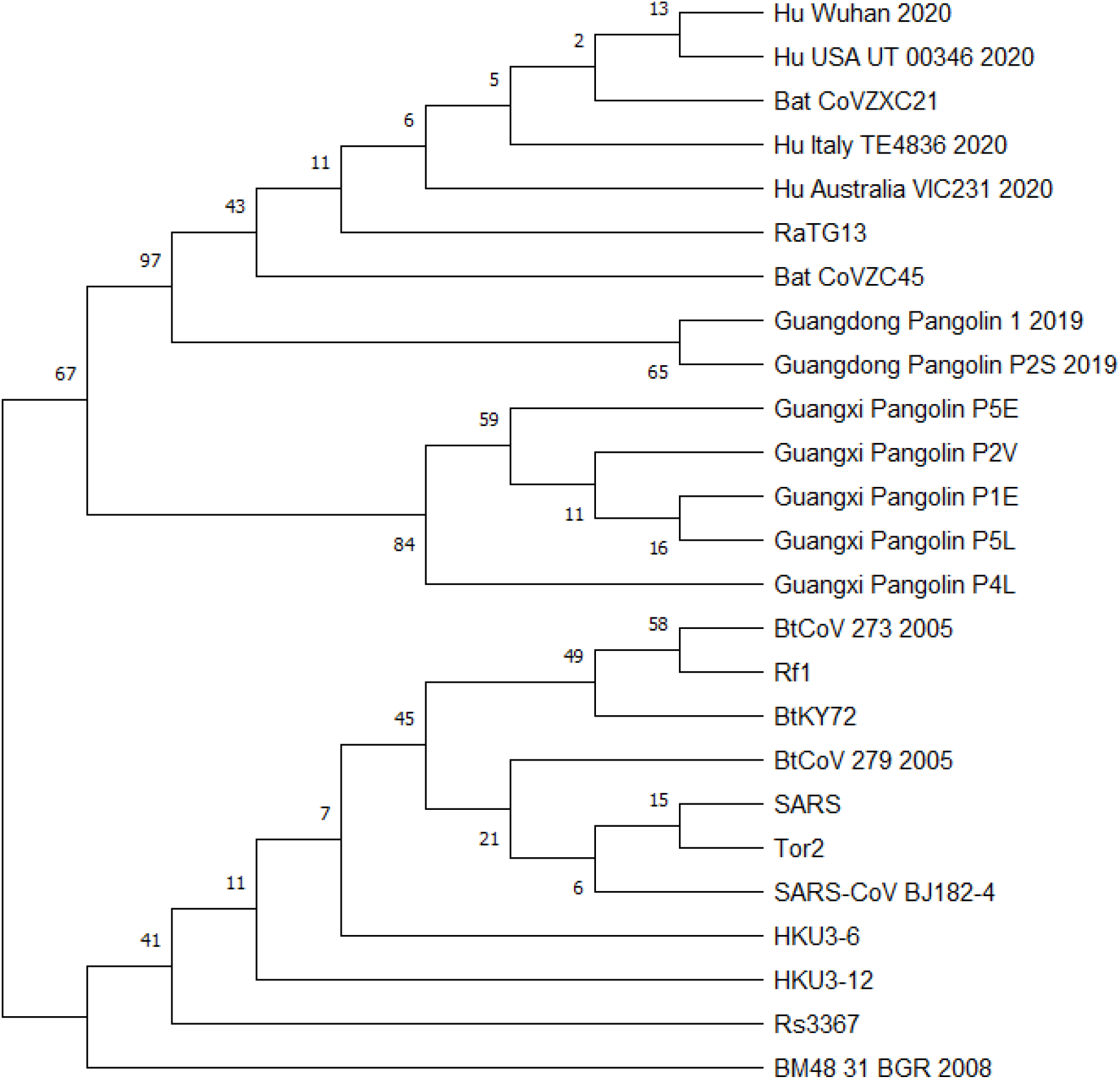
Phylogenetic tree of gene ORF10 inferred using RAxML with 100 bootstrap replicates for a group of 25 betacoronaviruses (see Supplementary Table 1).

### Tree for RB domain (43 species)

**Supplementary Figure 12.**
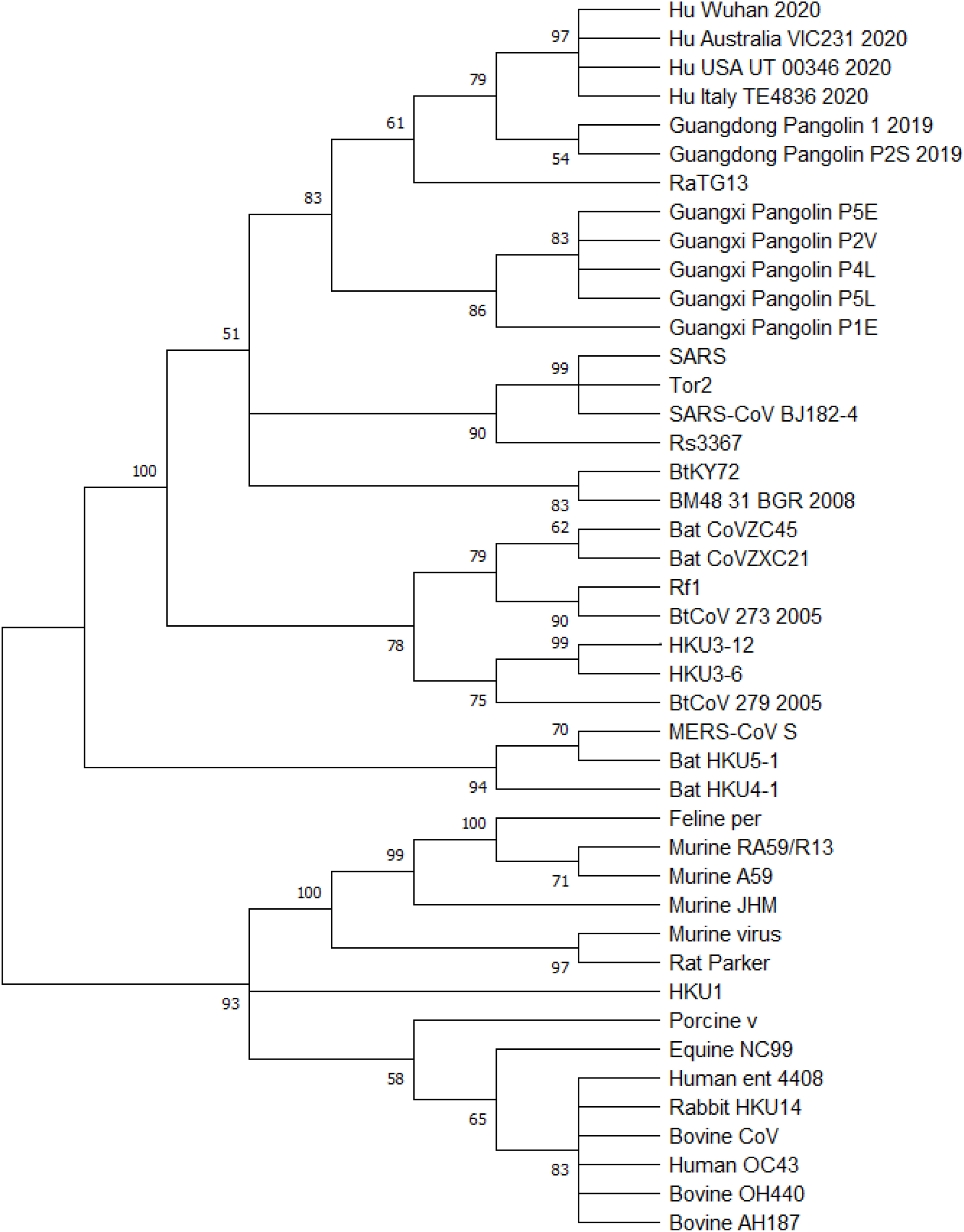
Phylogenetic tree of the RB domain inferred using RAxML with 100 bootstrap replicates for a group of 43 betacoronaviruses (see Supplementary Table 1).

